# Genomic potential for photoferrotrophy in a seasonally anoxic Boreal Shield lake

**DOI:** 10.1101/653014

**Authors:** JM Tsuji, N Tran, SL Schiff, JJ Venkiteswaran, LA Molot, M Tank, S Hanada, JD Neufeld

## Abstract

Photoferrotrophy, the light-induced oxidation of ferrous iron, is thought to have contributed to primary production within Earth’s early anoxic oceans yet is presumed to be of little modern environmental relevance. Here we use genome-resolved metagenomics and enrichment cultivation to explore the potential for photoferrotrophy in the anoxic water columns of globally abundant Boreal Shield lakes. We recovered four high-completeness and low-contamination draft genome bins assigned to the class *Chlorobia* (formerly phylum *Chlorobi*) from environmental metagenome data and enriched two novel sulfide-oxidizing species, also from the *Chlorobia*. The sequenced genomes of both enriched species, including the novel “*Candidatus* Chlorobium canadense”, encoded the *cyc2* candidate gene marker for iron oxidation, suggesting the potential for photoferrotrophic growth. Surprisingly, one of the environmental genome bins encoded *cyc2* and lacked sulfur oxidation gene pathways altogether. Despite the presence of *cyc2* in the corresponding draft genome, we were unable to induce photoferrotrophy in “*Ca.* Chlorobium canadense”, suggesting that yet-unexplored mechanisms regulate expression of sulfide and ferrous iron oxidation gene systems, or that previously unrecognized functions for this outer membrane cytochrome exist. Doubling the known diversity of *Chlorobia*-associated *cyc2* genes, metagenome data showed that putative photoferrotrophic populations occurred in one lake but that only sulfide-oxidizing populations were present in a neighboring lake, implying that strong ecological or geochemical controls govern the favourability of photoferrotrophy in aquatic environments. These results indicate that anoxygenic photoautotrophs in Boreal Shield lakes could have unexplored metabolic diversity that is controlled by ecological and biogeochemical drivers pertinent to understanding Earth’s early microbial communities.

## Introduction

Anoxygenic photoautotrophs are common in oxygen-limited environments worldwide and serve as primary producers in sediments, hot springs, and anoxic water columns as deep as the Black Sea [1–3]. Microorganisms performing anoxygenic photosynthesis commonly use a reduced sulfur compound, such as sulfide or thiosulfate, as the photosynthetic electron donor [4, 5]. However, the discovery of alternative metabolic modes of photosynthesis challenge classical interpretations of how anoxic ecosystems function [6–8]. Photoferrotrophy, an anoxygenic photosynthetic process involving the oxidation of ferrous iron in place of reduced sulfur, has been described in several laboratory cultures but remains poorly understood with respect to its ecological role [9, 10]. Photoferrotrophs are of great interest for their potential involvement in early Earth microbial communities where they may have served as important primary producers in the ferruginous (i.e., iron-rich, sulfate-poor) and anoxic waters of Archaean Eon oceans (ca. 3.8-2.5 Ga) [11–13]. However, photoferrotrophy remains poorly explored in modern environments, and the process is assumed to have little ecological importance given the oxic nature of the Earth’s atmosphere and higher levels of sulfate in most modern aquatic systems compared to Archaean oceans [10].

One of the key challenges in assessing the modern importance of photoferrotrophy has been a limited understanding of the biochemistry and genetics of iron oxidation. However, recent work has begun to unravel the molecular basis for iron oxidation in photoferrotrophs and other neutrophilic iron-oxidizing bacteria (reviewed in [14, 15]). Candidate gene systems are emerging that can be used to probe for selected types of iron oxidation environmentally. In particular, genes encoding for porin-cytochrome *c* protein complexes (PCC), such as *pioAB* and *mtoAB*, and genes encoding for outer-membrane monoheme c-type cytochromes, such as *cyc2*, have been identified as potentially useful markers for iron oxidation [15–19]. Recent environmental metatranscriptomic work, for example, suggests tight links between zetaproteobacterial *cyc2* gene expression and iron oxidation for these microaerophilic iron-oxidizing bacteria [20]. The discovery of such genetic markers expands capabilities to explore the role of photoferrotrophy and other forms of iron cycling in nature.

Novel photoferrotrophic bacteria are also being isolated in the laboratory. For example, among the *Chlorobia* class (i.e., “green sulfur bacteria” and “*Candidatus* Thermochlorobacter aerophilum”), the only known photoferrotroph was *Chlorobium ferrooxidans* strain KoFox, which was enriched from freshwater lake sediments [21]. Recently, *Chlorobium phaeoferrooxidans* strain KB01, a bacteriochlorophyll *e*-containing member of the class *Chlorobia*, was isolated from the anoxic water column of the meromictic and ferruginous Kabuno Bay [22]. In addition, *Chlorobium* sp. strain N1 was isolated from marine sediments [23]. Both *Chl. ferrooxidans* and *Chl. phaeoferrooxidans* oxidize ferrous iron as their sole photosynthetic electron donor, assimilating sulfur as sulfate [21, 22]. *Chl.* sp. N1 is capable of oxidizing either ferrous iron or sulfide as the photosynthetic electron donor and can also utilize organic compounds [23]. A fourth recently isolated photoferrotroph, *Chlorobium* sp. BLA1, was shown to also be capable of oxidizing sulfide, although genome data is not yet available for this strain [24]. Phylogenetically, *Chl. ferrooxidans* and *Chl. phaeoferrooxidans* are sister groups to one another, whereas *Chl.* sp. N1 is closely related to *Chl. luteolum*, a sulfide-oxidizing species of *Chlorobia* hypothesized to be able to switch to photoferrotrophic growth [22, 23, 25]. Genome sequences available for *Chl. ferrooxidans, Chl. phaeoferrooxidans, Chl.* sp. N1, and *Chl. luteolum* show that their genomes encode *cyc2* homologues, unlike all other sulfide-oxidizing *Chlorobia* [25–28]. These recent findings allow for first glimpses into the natural diversity of photoferrotrophs and provide some evidence that *cyc2* may serve as a genomic marker for photoferrotrophy within the class *Chlorobia*.

Current knowledge of photoferrotroph ecology derives from studies of a select few modern environments. Among the best studied are meromictic ferruginous lakes that are used as analogues of Archaean ocean conditions (reviewed in [29]; see also [30, 31]). Study of these Archaean ocean analogues has led to varying conclusions about the environmental importance of photoferrotrophy. In Lake Matano (Indonesia), bacteria classified to the *Chlorobium* genus were detected beneath the oxic-anoxic zone interface [32]. However, only genes implicated in the sulfide oxidation pathway among *Chlorobia* were detected in metagenome sequencing data, and anoxygenic photoautotrophy rates were below detection limits, leaving the role of photoferrotrophy unclear [33]. In Lake La Cruz (Spain), rates of photoferrotrophic iron oxidation were estimated at 0.2-1.4 µM Fe^2+^ d^−1^, yet microscopy data and photoautotrophy rates suggested that photoferrotrophy was catalyzed by a small proportion of total phototrophic cells, and that photoferrotrophs co-existed with other phototrophic bacteria, such as sulfide oxidizers [34]. In Lake Kivu’s Kabuno Bay (Democratic Republic of the Congo), anoxygenic phototrophs in the upper anoxic zone used ferrous iron for most or all photosynthetic activity at rates of up to 100 µM Fe^2+^ d^−1^ [22]. Anoxygenic photoautotrophs were estimated as being capable of supplying up to ∼90% of total fixed carbon in the bay’s anoxic zone [35]. Contrasting results from these lakes suggest that unknown factors in ferruginous systems modulate the relative importance of various anoxygenic photosynthetic modes. In addition, it appears as though sulfide-oxidizing photoautotrophs can play a relatively important role in primary production even in sulfide-limited systems (see also [36]). However, the limited number of such meromictic ferruginous lakes makes further speculation about photoferrotroph ecology and *in situ* activity challenging.

Recently, we combined 16S ribosomal RNA (16S rRNA) gene sequencing and iron isotope data to propose photoferrotrophy by populations of *Chlorobia* as an important photosynthetic process in the seasonally anoxic water columns of two ferruginous Boreal Shield lakes at the International Institute for Sustainable Development Experimental Lakes Area (IISD-ELA; near Kenora, Canada) [37]. We studied Lake 227, an experimentally eutrophied lake, and Lake 442, a nearby and pristine reference lake [38, 39]. Boreal Shield lakes number in the tens of millions across northern regions globally, and we proposed based on our preliminary data that Boreal Shield lakes that develop seasonal anoxia commonly contain photoferrotrophic bacteria and could serve as plentiful modern-day analogues of Archaean ocean ecosystems. Here, we use genome-resolved environmental metagenomics to recover high-completeness, low-contamination draft genomes of key populations of *Chlorobia* from the same two IISD-ELA lakes. We then apply recent knowledge of iron functional gene markers to test whether these populations possess the genetic potential for photoferrotrophy or other phototrophic modes. We also report the enrichment of two novel species of *Chlorobia* from Lake 227 and nearby Lake 304 and analyze their genetic and functional potential for phototrophic sulfide and iron oxidation. Through this first usage of molecular tools to explore *Chlorobia*-driven photoferrotrophy in an environmental context, we aim to clarify the diversity and ecology of anoxygenic photoautotrophic bacteria in Boreal Shield lakes, with implications for understanding both Archaean microbial communities and modern ferruginous environments.

## Materials and Methods

### Lake sampling, metagenome sequencing, assembly, and binning

The IISD-ELA sampling site, lake geochemistry, and sample collection methods were described in detail previously [37]. Based on preliminary 16S rRNA gene data, we selected six water column genomic DNA samples collected from Lakes 227 and 442 for re-sequencing via shotgun metagenomics. All six samples were collected at or beneath the oxic-anoxic zone boundary, at depths where low light penetration occurs, and had high relative abundances of anoxygenic phototrophs, dominated by populations of *Chlorobia*. Samples from Lake 227 were selected at 6 m and 8 m in 2013 and 2014, and samples from Lake 442 were selected at 16.5 m in 2011 and 13 m in 2014. The sequencing library was prepared using the NEBNext Ultra II DNA Library Prep Kit for Illumina (New England Biolabs; Ipswich, Massachusetts, USA) and was sequenced on a single lane of a HiSeq 2500 (Illumina; San Diego, California, USA) in Rapid Run Mode with 2×200 base paired-end reads, generating 26.3 to 39.5 million reads per sample. Library preparation and sequencing was performed by the McMaster Genome Facility (Hamilton, Ontario, Canada).

Raw metagenome reads were quality-trimmed, assembled, binned, and annotated using the ATLAS pipeline, version 1.0.22 [40]. The configuration file with all settings for the pipeline is available in Supplementary File 1. To improve genome bin completeness and reduce bin redundancy, QC processed reads from related samples (i.e., L227 2013 6 m and 8 m, L227 2014 6 m and 8 m, and L442 samples) were co-assembled and re-binned using differential abundance information via a simple wrapper around ATLAS, as described in the Supplementary Methods.

### Identification of cyc2 genes and Chlorobia genome bins

Assembled contigs were assessed for the presence of *cyc2* gene homologues by building a custom profile Hidden Markov Model (HMM). The predicted primary sequences of *cyc2* were recovered from the four available genomes of *Chlorobia* known to possess the gene, *Chl. ferrooxidans, Chl. phaeoferrooxidans, Chl.* sp. N1, and *Chl. luteolum*, as well as from the genomes of reference microaerophilic iron-oxidizing bacteria as described by He and colleagues [14]. A cytochrome 572 gene from *Leptospirillum* sp. (EDZ39515.1) was omitted from the reference dataset, due to its high divergence from other sequences, to build a more robust alignment for the *cyc2* clade relevant to the *Chlorobia* [41]. Collected sequences were aligned using Clustal Omega, version 1.2.3 [42], and the alignment was used to build a profile HMM using the *hmmbuild* function of HMMER3, version 3.2.1 [43]. Recovered *cyc2* genes over the course of the study were added to the alignment and HMM; final HMMs are included in Supplementary File 2.

Recovered *cyc2* genes were compared to *cyc2* reference sequences by building a maximum-likelihood phylogeny. The alignment was masked using Gblocks, version 0.91b [44], as described in the Supplementary Methods. The maximum likelihood phylogeny was then prepared from the masked sequence alignment via IQ-TREE, version 1.6.10 [45], using the LG+F+I+G4 sequence evolution model as determined by the ModelFinder module of IQ-TREE [46]. To build the consensus tree, 1000 bootstrap replicates were performed, each requiring ∼100-110 tree search iterations for phylogeny optimization.

Genome bins were dereplicated using dRep version 2.0.5 [47] with default settings. All genome bins from individual assembles and co-assemblies were pooled for dereplication to help ensure that the highest quality bins, whether from individual assemblies or co-assemblies, would be retained for downstream analyses. Genome bins of *Chlorobia* with > 90% completeness and < 10% contamination based on CheckM statistics [48] were selected for further study. These genome bins were imported into Anvi’o version 4 [49] and were manually examined for contigs improperly binned based on read mapping, tetranucleotide frequencies, and contig taxonomic classification. Import of ATLAS data into Anvi’o was performed using the *atlas-to-anvi.sh* script, version 1.0.22-coassembly-r4, available in the *atlas-extensions* GitHub repository at https://github.com/jmtsuji/atlas-extensions. Contigs in curated genome bins of *Chlorobia* were then ordered via the Mauve Contig Mover development snapshot 2015-02-13 for Linux [50] using the *Chl. luteolum* genome to guide ordering. To assess bin quality, tRNA genes were predicted using Prokka v1.13.3 [51].

### Enrichment cultivation, sequencing, and assembly

Lake 227 and Lake 442 were sampled again in June 2016 and July 2017, along with the nearby Lake 304 in July 2017. Basic sampling information for each lake is summarized in Supplementary File 3. Lake 304 has been described previously and also has seasonally anoxic bottom waters [52, 53]. The lake was experimentally fertilized with phosphate in 1971-72 and 1975-76, with ammonium in 1971-74, and with nitrate in 1973-74, but the lake has not been manipulated since then and rapidly returned to its non-eutrophied state once additions ceased, having a water residence time of ∼2.7 years [54–56]. Lake 304 was added to our sampling efforts for enrichment culturing to explore the broader distribution of photoferrotrophy among IISD-ELA lakes, because preliminary 16S rRNA gene sequencing data (not shown) indicated high relative abundances of *Chlorobia* populations in this lake. Anoxic water was collected from Lakes 227 and 304 at a depth of 6 m and Lake 442 at 15 m, where trace levels of light are generally detectable (i.e., in the range of 0.01-1 µmol photons m^−2^ s^−1^ between 400-700 nm wavelengths, as commonly measured when field sampling). Lake water was pumped to the surface using a gear pump and directly injected into N_2_-filled 160 mL glass serum bottles that were sealed with blue butyl rubber stoppers (Bellco Glass; Vineland, New Jersey, USA), using a secondary needle to vent N_2_ gas until the bottles were full. Water was kept cold (∼4°C) and dark after collection and during shipping.

Enrichment cultures were grown using sulfide-containing Pfennig’s medium, prepared as described by Imhoff [4, 5], or ferrous iron-containing freshwater medium, prepared as described by Hegler and colleagues [57]. The ferrous iron-containing medium contained 8 mM ferrous chloride (FeCl_2_), without filtration of precipitates, and used trace element solution SLA [4]. Initial enrichments contained 10-20% lake water at a total volume of 50 mL and were inoculated into 120-160 mL glass serum bottles sealed with blue butyl rubber stoppers (Bellco Glass) or black rubber stoppers (Geo-Microbial Technology Company; Ochelata, Oklahoma, USA). The remaining headspace was flushed with a 90:10 N_2_:CO_2_ gas mix at left at a pressure of 1.5 atm. Several bottles additionally had anoxic DCMU (i.e., Diuron or 3-(3,4-dichlorophenyl)-1,1-dimethylurea; Sigma-Aldrich; St. Louis, Missouri, USA) added to a concentration of 50 µM to block activity of oxygenic phototrophs [58, 59]. Bottles were incubated at 22°C and ∼50 cm away from far red PARSource PowerPAR LED Bulbs (LED Grow Lights Depot; Portland, Oregon, USA) as the light source.

Two enrichment cultures survived repeated subculturing or dilution-to-extinction and contained green sulfur bacteria based on pigmentation and marker gene sequence analysis. Biomass from these cultures was collected by centrifugation, and genomic DNA was extracted using the DNeasy UltraClean Microbial Kit (Qiagen; Venlo, The Netherlands). Genomic DNA was prepared into metagenome sequencing libraries using the Nextera DNA Flex Library Prep Kit (Illumina), and the library was sequenced on a fraction of a lane of a HiSeq 2500 (Illumina) in High Output Run Mode with 2×125 base paired-end reads. Library preparation and sequencing was performed by The Centre for Applied Genomics (TCAG; The Hospital for Sick Children, Toronto, Canada), generating ∼6 million total reads per sample. Metagenomes were assembled using ATLAS version 1.0.22 without co-assembly. Genome bins of *Chlorobia* were manually refined as described above, using Anvi’o version 5 [49] via *atlas-to-anvi.sh* commit 99b85ac; contigs of curated bins were subsequently ordered, and tRNA genes counted, as described above.

### Comparative genomics of Chlorobia genomes

Refined genome bins of *Chlorobia*, which belonged to the *Chlorobium* genus, were compared to genomes of reference strains from the *Chlorobiaceae* family. Genomes of all available type strains and photoferrotrophs from the family were downloaded from the NCBI (Supplementary File 4). Average nucleotide identity (ANI) between genomes was calculated using FastANI version 1.1 [60]. The phylogenetic relationship between genomes was determined by constructing a concatenated ribosomal protein alignment based on the rp1 set of 16 ribosomal protein genes using GToTree version 1.1.10 [61, 62]. IQ-TREE version 1.6.9 [45] was used to construct the maximum likelihood phylogeny from the alignment. The LG+F+R4 model of sequence evolution was identified as optimal by the IQ-TREE ModelFinder module [33], and phylogeny construction used 1000 bootstrap replicates, each requiring ∼100-110 tree search iterations for optimization.

The presence or absence of genes implicated in iron or sulfur cycling metabolism in the collection of *Chlorobiaceae* genomes was assessed using reciprocal BLASTP [63]. Genes were selected from the genomes of *Chl. ferrooxidans* and *Chl. clathratiforme* based on the genome comparison of Frigaard and Bryant [25]. Predicted primary sequences of these genes were used to identify putative homologues across other genomes of *Chlorobia* using the BackBLAST pipeline, version 2.0.0-alpha2 (doi:<u>10.5281/zenodo.3465955</u>) [64]. The e-value cutoff for BLAST hits was set at 10^−40^, based on empirical testing of gene clusters known to be present or absent in reference genomes of *Chlorobia*, and the identity cutoff was set to 20%. Selected genes (e.g., *qmoA, dsrJ*) were omitted due to poor homology between reference genomes. Genes associated with photosynthesis and carbon fixation were similarly assessed using BackBLAST, except that most reference genes were selected from the genome of *Chl. tepidum* according to Bryant and colleagues [65] and Tourova and colleagues [66].

### Metagenome taxonomic and functional profiling

Environmental relative abundances of microbial populations were estimated by read mapping to dereplicated genome bins (further details in Supplementary Methods). Taxonomy and functional gene information were then overlaid onto relative abundance data. Genome bin taxonomy was determined based on the Genome Taxonomy Database (GTDB) using GTDB-Tk, version v0.2.2, relying on GTDB release 86, version 3 (April 2019) [67, 68]. As such, GTDB taxonomy names are used throughout this manuscript. Genomes were tested for the presence of the *cyc2* and *dsrA* functional genes using the HMM developed in this study and an HMM available on FunGene [69], respectively. Gene amino acid translations were predicted for all genome bins using prodigal version 2.6.3 [70], via the GTDB-Tk, and amino acid translations were searched via HMMs using the MetAnnotate pipeline development release version 0.9.2 [71]. An e-value cutoff of 10^−40^ was used to filter low-quality gene hits. In case of bias due to unassembled or incorrectly binned genes, genome bin-based environmental abundance metrics were cross-compared to abundance metrics generated by directly scanning unassembled metagenome reads, as described in the Supplementary Methods.

### Assessment of ferrous iron oxidation potential of Chlorobia enrichments

After initial sequencing, cultures containing *Chlorobia* spp. continued to be purified in the laboratory. A single culture, which was enriched on sulfide-containing medium, survived continued cultivation and was provisionally named “*Candidatus* Chlorobium canadense strain L304-6D” (ca.na.den’se N.L. neut. adj. *canadense* from or belonging to Canada). “*Ca*. Chl. canadense” was purified through multiple rounds of incubation in deep agar dilution series and picking of isolated green colonies [72]. For deep agar shakes, Pfennig’s medium or modified *Chloracidobacterium thermophilum* Midnight (CTM) medium [73] containing 1-3 mM buffered sulfide feeding solution were used [4]. Eventually, in preparation for growth in ferrous iron-containing medium, cultures of “*Ca.* Chl. canadense” were transferred to liquid freshwater medium as described by Hegler and colleagues [57] (here termed, “Hegler freshwater medium”) but containing 0.5-1 mM buffered sulfide feeding solution, 10-20 µM FeCl_2_, and 0.5 mg/L resazurin to monitor the medium’s redox status.

To test its ability to oxidize ferrous iron phototrophically, “*Ca.* Chl. canadense” was inoculated into Hegler freshwater medium containing low levels of ferrous iron. Hegler freshwater medium was made without ferrous iron nor sulfide and was aliquoted into 120 mL glass serum bottles under flow of sterile 90:10 N_2_:CO_2_ gas, with 50 mL of medium per bottle. Bottles were sealed with black rubber stoppers (Geo-Microbial Technology Company) and were bubbled for at least nine minutes with additional 90:10 N_2_:CO_2_ gas to reduce trace oxygen. The headspace of each bottle was left at 1.5 atm pressure. A sterile solution of FeCl_2_ was added to each bottle to reach a ferrous iron content of 100 μM, and to some bottles, a sterile and anoxic solution of the chelator ethylenediaminetetraacetic acid (EDTA) was added to a concentration of 120 μM as done by Peng and colleagues [74] to explore whether this chelator could enhance photoferrotrophic iron oxidation. Bottles were incubated at room temperature in the dark overnight to allow for complexation of ferrous iron and EDTA. All media was confirmed to have a pH of 6.5-7.

“*Ca.* Chl. canadense” was grown in 100 mL Hegler freshwater medium using a total of ∼2 mM sulfide, fed incrementally in 0.4 mM doses, until sulfide was completely oxidized and the resazurin in the bottle went slightly pink. At the same time, a culture of *Chl. ferrooxidans* KoFox was grown in the same freshwater medium containing ∼8 mM FeCl_2_ until ferrous iron was completely oxidized. The entire 100 mL “*Ca.* Chl. canadense” culture was pelleted by centrifugation at 7000 x g for 13 minutes, and the pellet was washed twice with unamended Hegler freshwater medium. The ∼200 mg of wet biomass recovered was suspended in 1 mL of unamended freshwater medium, and 0.1 mL was inoculated into each relevant incubation bottle. Similarly, 100 mL of the *Chl. ferrooxidans* reference culture was pelleted by centrifugation at 7000 x g for 5 minutes, washed twice, and suspended in 1 mL of unamended freshwater media, and 0.1 mL was added to relevant bottles. Only the surfaces of *Chl. ferrooxidans* cell pellets, which contained mostly green cells and not brown ferric iron precipitate, were suspended with each wash, allowing for ∼3 mg of nearly iron-free wet biomass to be recovered. Once cultures were inoculated, they were transferred to a 22°C incubator without shaking where they received 30 μmol photons m^−2^ s^−1^ white light from a mix of fluorescent (F48T12/CW/VHO; Osram Sylvania; Wilmington, Massachusetts, USA) and incandescent (60W) bulbs.

Cultures were sampled regularly over a 21-day incubation period to monitor iron concentrations and iron oxidation states. At each sampling time point, an aliquot of culture was removed from each bottle using a sterile 90:10 N_2_:CO_2_-flushed syringe and 25 G needle, and 330 μL of this culture aliquot was acidified immediately with 30 μL of 6 N hydrochloric acid (HCl). Acidified samples were stored for no more than two days at 4°C (with the exception of samples collected on day 14; see Supplementary Methods) before being assessed for iron species concentrations via the ferrozine assay [75], with 5% ascorbic acid being used as the reducing agent to determine total iron species [76].

## Results

### Recovery of Chlorobium genome bins

Binning of assembled contigs from the lake and enrichment culture metagenomes, followed by dereplication of highly similar bins and manual curation, allowed for the recovery of six highly complete, low-contamination genome bins that classified within the genus *Chlorobium* (Table 1). Three of the genome bins recovered from lake metagenomes had best representatives selected by dRep from the Lake 227 (2013 sample) co-assembly, and one had its best representative from the Lake 442 (2011/2014 sample) co-assembly. In addition, a single *Chlorobium* genome bin was recovered from each enrichment culture. Recovered bins had at least 90.1% completeness and a maximum of 2.8% contamination based on analysis by CheckM [48]. Recovered genome bins were between 1.9 to 2.6 Mb in size, within the approximate length range of reference genomes associated with the family *Chlorobiaceae* (2.0-3.3 Mb; average 2.7 Mb), and were represented by 13 to 208 contigs. Other lower quality genome bins classified as belonging to the *Chlorobia* were also recovered (Supplementary File 5), but these were not considered for gene pathway analysis in order to minimize the risk of including false positives. In particular, L442 Bin 74 was excluded from the set of curated bins due to its high contig count (282) and elevated predicted contamination (4.12%) and strain heterogeneity (25%).

The constructed HMM for the *cyc2* gene enabled the recovery of five potential *cyc2* homologues associated with *Chlorobia* from assembled contigs (Fig 1). Three of these *cyc2* homologues were found in the set of six manually curated genome bins described above, and another homologue was detected in one of the lower quality genome bins of *Chlorobia* (L227 2014 Bin 92). The final homologue was detected on a short contig in the same lower quality genome bin and was removed from subsequent analyses due to its lack of gene neighbours. Detected *cyc2* homologues were full-length genes, and their predicted primary sequences were detected in the HMM search with a maximum e-value of ∼10^−140^. All recovered *cyc2* homologues contained the heme-binding CXXCH motif near the N-terminus of their predicted primary sequence, and all homologues were predicted by I-TASSER [77] to contain a porin-like beta barrel structure for membrane binding (Fig. 1A; one example shown) [13]. Aside from the closely related genomes of *Chl. ferrooxidans* and *Chl. phaeoferrooxidans* (Supplementary Figure 1), all other genomes showed substantial genome re-arrangement around the *cyc2* homologue (Fig. 1B). However, detected *cyc2* homologues were always adjacent to a *c5* family cytochrome (Fig. 1B). These *c5* family cytochromes had low sequence identity to one another (i.e., as low as 30%) yet also contained a CXXCH motif near the N-terminal end (Supplementary File 6). All recovered *cyc2* homologues grouped monophyletically with *cyc2* predicted primary sequences belonging to reference genomes of *Chlorobia* compared to the *cyc2* of other reference strains (Fig. 1C). Overall, the combination of HMM search specificity, genomic context, predicted gene motifs, and phylogenetic placement is strong evidence that the identified cytochrome genes are *cyc2* homologues.

**Figure 1.**
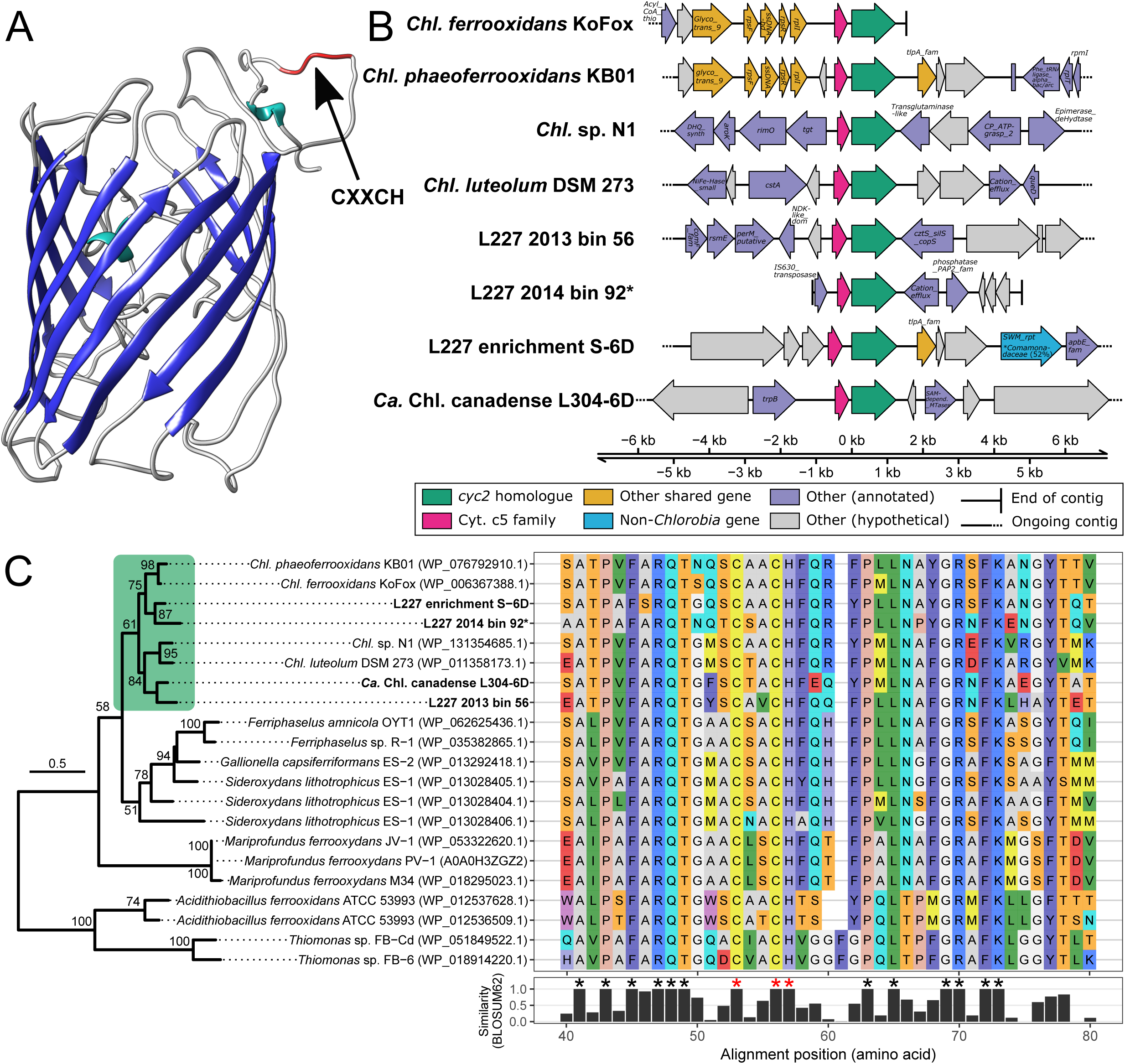
Recovered *cyc2* homologues of *Chlorobia* from metagenomes. **A**, Homology model of the Cyc2 protein from *Chl. phaeoferrooxidans* KB01, generated by I-TASSER based on sequence homology (with leading 18 signal peptides trimmed according to SignalP [94]) and visualized using UCSF Chimera [95]. The porin-like beta barrel structure is highlighted in dark blue, and the location of the single N-terminal heme-binding motif on the outer membrane portion of the structure is indicated by a black arrow. **B**, Genomic context of the recovered *cyc2* homologues. A ∼10 kb region is shown of assembled contigs surrounding the homologues. The *cyc2* and adjacent c5 family cytochrome genes are highlighted in green and pink, respectively. Other shared genes between contigs are highlighted in yellow. Remaining genes are coloured grey if they lacked a functional annotation (e.g., no significant BLASTP hit, or hit to a domain of unknown function) or are coloured purple if their closest BLASTP hit corresponded to a gene associated with the *Chlorobiaceae* family. One gene in the 10 kb region had its closest BLASTP hit to a gene associated with the *Comamonadaceae* family (*Betaproteobacteria*; ∼51% amino acid identity); this gene is highlighted in blue. **C**, Sequence comparison of recovered *cyc2* homologues to *cyc2* genes of known iron-oxidizing microorganisms. The phylogenetic tree was built from a 223 residue masked Cyc2 amino acid sequence alignment (see Methods). Bootstrap values over 50/100 are shown, and the scale bar represents the proportion of residue changes along the alignment. Due to the uncertain evolutionary history of *cyc2*, the phylogeny is midpoint-rooted. A green box highlights the monophyletic *Chlorobia* clade. Adjacent to the phylogeny, a subset of the unmasked Cyc2 sequence alignment is shown (positions 40-80 of 609; N-terminus end). Alignment positions having 100% sequence identity in the predicted heme-binding site (CXXCH motif) are marked with red asterisks. Other positions with 100% sequence identity are marked with black asterisks. The full sequence alignment is available in Supplementary File 6. Note that L227 2014 Bin 92 (name marked with an asterisk) was included in this figure for comparison of its *cyc2* gene despite the bin quality being lower than the main *Chlorobia* genome bin set. Abbreviations: *Chl.* = *Chlorobium*.

### Enrichment cultivation of Chlorobia

Enrichment cultivation was attempted with anoxic lake water using both sulfide- and ferrous iron-containing anoxic media. For Lake 442, one enrichment in a ferrous iron-containing medium grew with evidence of light-driven iron oxidation, but 16S rRNA gene sequencing showed that the culture was dominated by a member of the genus *Rhodopseudomonas*, which did not represent a detectable genus in environmental sequencing data. Given the known metabolic versatility of *Rhodopseudomonas* spp. and the negligible contribution of members of this genus to the studied lake microbial communities, this culture was not pursued further. No members of the *Chlorobia* were ever detected in any enrichment culture from Lake 442.

For Lakes 227 and 304, growth was observed for both media tested. Enrichments grown on sulfide-containing media developed green colour after ∼12-15 weeks of incubation and contained populations of *Chlorobia* based on Sanger sequencing of both the V3-V4 region of the 16S rRNA gene and the partial *dsrAB* gene (using *Chlorobia*-targeted PCR primers 341f/GSB822r [78] and PGdsrAF/PGdsrAR [79], respectively). However, the appearances of cultures grown on ferrous iron-containing media for these lakes differed from those of reference photoferrotrophic members of *Chlorobia*. Instead of forming a reddish-brown ferric iron precipitate, the enrichments blackened, potentially indicative of sulfate reduction and metal sulfide formation. After blackening, enrichments developed a green colour (Supplementary Figure 2). Suspecting that sulfate reduction was occurring in the ferrous iron enrichment bottles to support sulfide-oxidizing phototrophy, one Lake 227 enrichment was transferred into sulfide-containing media and still developed green pigmentation, albeit slowly (Supplementary Figure 2). Amplification and Sanger sequencing of the partial *dsrAB* gene cluster from this enrichment showed that the gene sequence only differed by one ambiguous base from, and thus was essentially identical to, a corresponding Lake 227 sulfide enrichment. Ferrous iron enrichments eventually stopped being followed due to long growth times, low biomass, and instability of the cultures. Enrichments on sulfide-containing media were continued instead.

Metagenome sequencing of the two successful enrichments of *Chlorobia* grown on sulfide (L227 enrichment S-6D and L304 enrichment S-6D) showed that that genomes bins of the *Chlorobia* from both enrichments encoded *cyc2* homologues. Although L227 enrichment S-6D ceased to grow in laboratory culture, the L304 enrichment S-6D continued to be purified and was named “*Candidatus* Chlorobium canadense strain L304-6D”. At the time of running the ferrous iron oxidation test (below), the “*Ca.* Chl. canadense” culture consisted of >90 % cells of *Chlorobia* based on epifluorescence microscopy, and other contaminating cells were known to be chemoorganoheterotrophs and sulfate reducing bacteria based on previous 16S rRNA gene amplicon sequencing data (Supplementary File 7).

### Metabolic diversity and phylogeny of Chlorobium genome bins

Recovered genome bins of *Chlorobia* were metabolically diverse and grouped into four general categories with respect to the presence of genes involved in iron and sulfur cycling (Fig. 2). One of the lake-recovered bins (L227 2013 Bin 56) lacked all genes involved in sulfide, elemental sulfur, and thiosulfate oxidation but contained the *cyc2* gene homologue. Conversely, two other lake-recovered bins (L227 2013 Bin 55 and L442 Bin 64) contained all assessed genes in the sulfide oxidation pathway but lacked the *cyc2* gene. The final lake-recovered bin (L227 2013 Bin 22) lacked both *cyc2* and genes for sulfur oxidation. This lack of key metabolism genes in L227 2013 Bin 22 might be due to incomplete genome binning, although the presence a bacterioferritin hypothesized by Frigaard and Bryant [25] to play a role in photoferrotrophy in this genome implies that photoferrotrophy might be possible. Lastly, the two enrichment culture bins contained genes involved in both sulfide and iron oxidation, similarly to *Chl.* sp. N1 and *Chl. luteolum*. Genes for hydrogen oxidation were detected in all of these genome bins (Fig. 2), along with marker genes for photosynthesis and bacteriochlorophyll biosynthesis (Supplementary Figure 3). Marker genes for carbon fixation (via the reverse TCA cycle) were detected in all genome bins, with the exception of L227 2013 Bin 56, which was missing one of the two assessed gene clusters for the this process (*aclAB*; Supplementary Figure 3).

**Figure 2.**
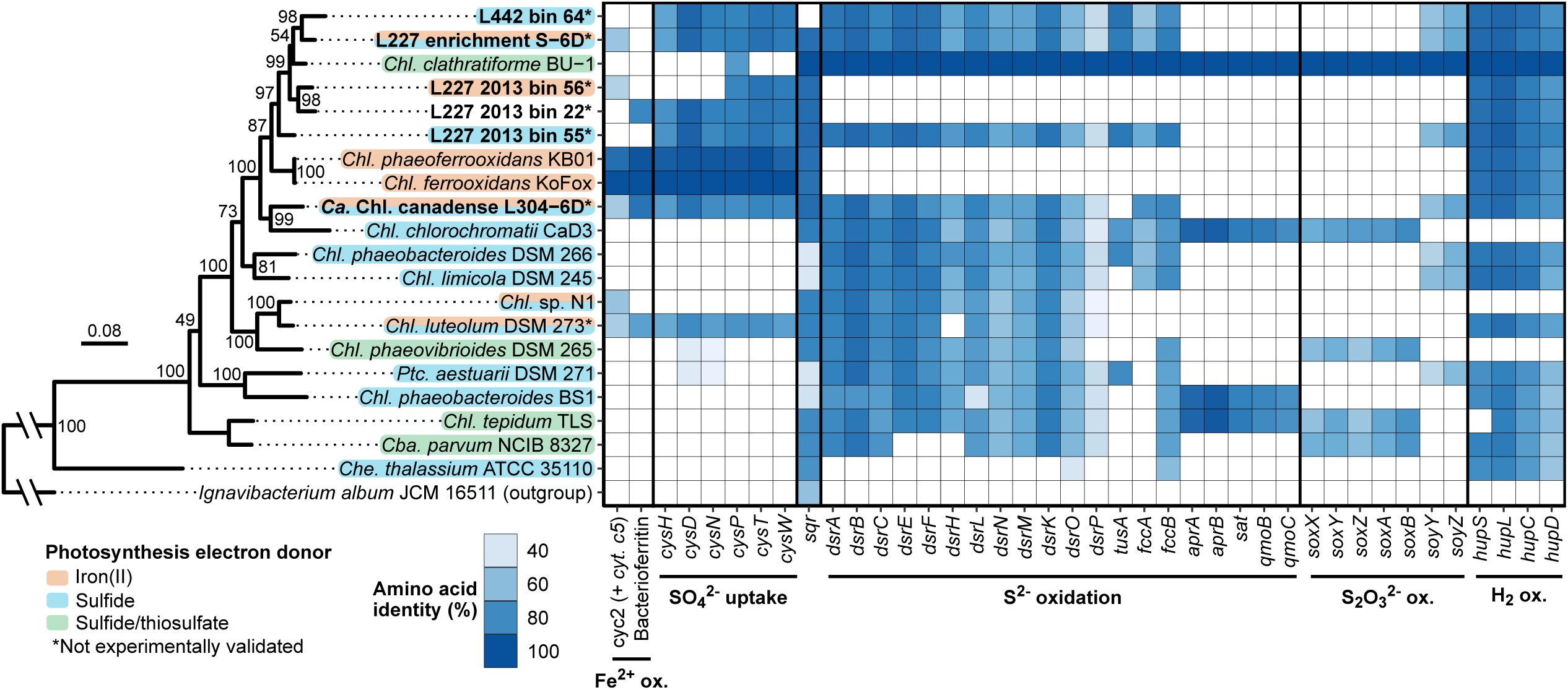
Iron- and sulfur-oxidizing genetic potential in recovered genome bins of *Chlorobia* compared to reference strains. The left side of the figure shows a maximum-likelihood phylogenetic tree of the *Chlorobia* based on concatenated ribosomal protein amino acid sequences (see methods). Bootstrap values over 50/100 are shown, and the scale bar represents the percentage of residue changes across the 2243 amino acid alignment. Species of *Chlorobia* are shaded based on their known or hypothesized metabolic potential. Genome bins of *Chlorobia* recovered from this study are bolded. On the right side, a heatmap is shown displaying the presence/absence of genes implicated in iron and sulfur metabolism among *Chlorobia* based on reciprocal BLASTP. Heatmap tiles are shaded based on the percent amino acid identity of an identified gene compared to the reference sequence (*Chl. ferrooxidans* for iron-related genes, and *Chl. phaeoclathratiforme* for sulfur-related genes). Although the cytochrome c5 family gene upstream of *cyc2* was not searched for directly due to poor sequence homology, the gene was verified manually to be adjacent to all hits of *cyc2*. Abbreviations: *Chl.* = *Chlorobium*; *Cba.* = *Chlorobaculum*; *Che.* = *Chloroherpeton*; *Ptc.* = *Prosthecochloris*.

All recovered *Chlorobia* genome bins encoded the *cys* gene cluster for assimilatory sulfate uptake, which was noted previously to potentially be associated with photoferrotrophy due to its absence from the genomes of most sulfur oxidizing strains [25]. The L227 2013 Bin 56 only contained half of the *cys* cluster, but the specific contig containing the cluster was checked and was found to have an assembly break point halfway through the gene cluster, making it likely that the complete *cys* cluster was present in the genome but failed to bin. However, the fact that *Chlorobia*-associated *cyc2* was completely undetectable in Lake 442 metagenome data (see below) implies that, although Lake 442 Bin 64 contains the *cys* cluster, it does not encode *cyc2*. Also, the genome of *Chl.* sp. N1 lacks the *cys* cluster, implying that the cluster is not necessary for photoferrotrophic growth (Fig. 2). As such, the *cys* cluster is likely not a reliable indicator of whether genomes of *Chlorobia* encode *cyc2*. Two of the genome bins also encoded putative homologues to a bacterioferritin that is potentially involved in photoferrotrophy (see above).

Phylogenetic analysis of the recovered *Chlorobium* genome bins based on concatenated ribosomal protein sequences showed that all four lake-recovered bins and the bin from the Lake 227 enrichment formed a monophyletic subgroup within the *Chlorobiaceae*, including *Chl. phaeoclathratiforme* BU-1 (Fig. 2). Directly basal to this subgroup was a branch containing *Chl. ferrooxidans* and *Chl. phaeoferrooxidans*. By comparison, the bin of “*Ca.* Chl. canadense” grouped sister to *Chl. chlorochromatii*, separately from the above group and from the group containing *Chl. luteolum* and *Chl.* sp. strain N1. Comparison of the concatenated ribosomal protein phylogeny to the *cyc2* predicted primary sequence phylogeny reveals some congruency between the two phylogenies for potentially photoferrotrophic members of the *Chlorobia*, with the exception of L227-S-6D and “*Ca*. Chl. canadense” (Supplementary Figure 4). Overall, all six recovered *Chlorobium* genome bins appear to represent novel species, having an ANI of < 79.6% compared to the genomes of all available type strains (Supplementary Figure 1).

### Functional and taxonomic profiling

Read mapping to the genome bins of *Chlorobia* showed that populations with varying genomic potential for iron and sulfur-cycling were relevant to the lake environment (Fig. 3). Bins containing *cyc2* (no *dsrA*) and bins containing *dsrA* (no *cyc2*) were both found at similar relative abundances in Lake 227 samples in 2013 and 2014 (Fig. 3A). Generally, genome bins of *Chlorobia* containing *cyc2* accounted for 0.6-2.8% of the Lake 227 microbial community data in the assessed samples, whereas bins of *Chlorobia* containing *dsrA* accounted for 1.7-4.1% of the data. Analyses of unassembled metagenome read data indicated similar relative abundances (Supplementary Figure 5). In contrast, only genome bins encoding sulfide oxidation were detected in Lake 442 metagenome assemblies. Even when unassembled metagenome data were scanned, only a single HMM hit from the Lake 442 metagenomes matched *cyc2* classified as belonging to a member of *Chlorobia*, compared to 27 000-37 000 total *rpoB* hits for these same metagenomes, showing that the lack of *cyc2* genes detected in genome bins of *Chlorobia* from L442 was not a result of incomplete assembly or genome binning. One genome bin of *Chlorobia* overlapped between L227 and L442 (L442 2011 16.5m Bin 8); the rest were distinct between the two lakes. In addition, the bins of *Chlorobia* recovered from the enrichment culture metagenomes were not detected at high relative abundances in any of the six lake metagenomes.

**Figure 3.**
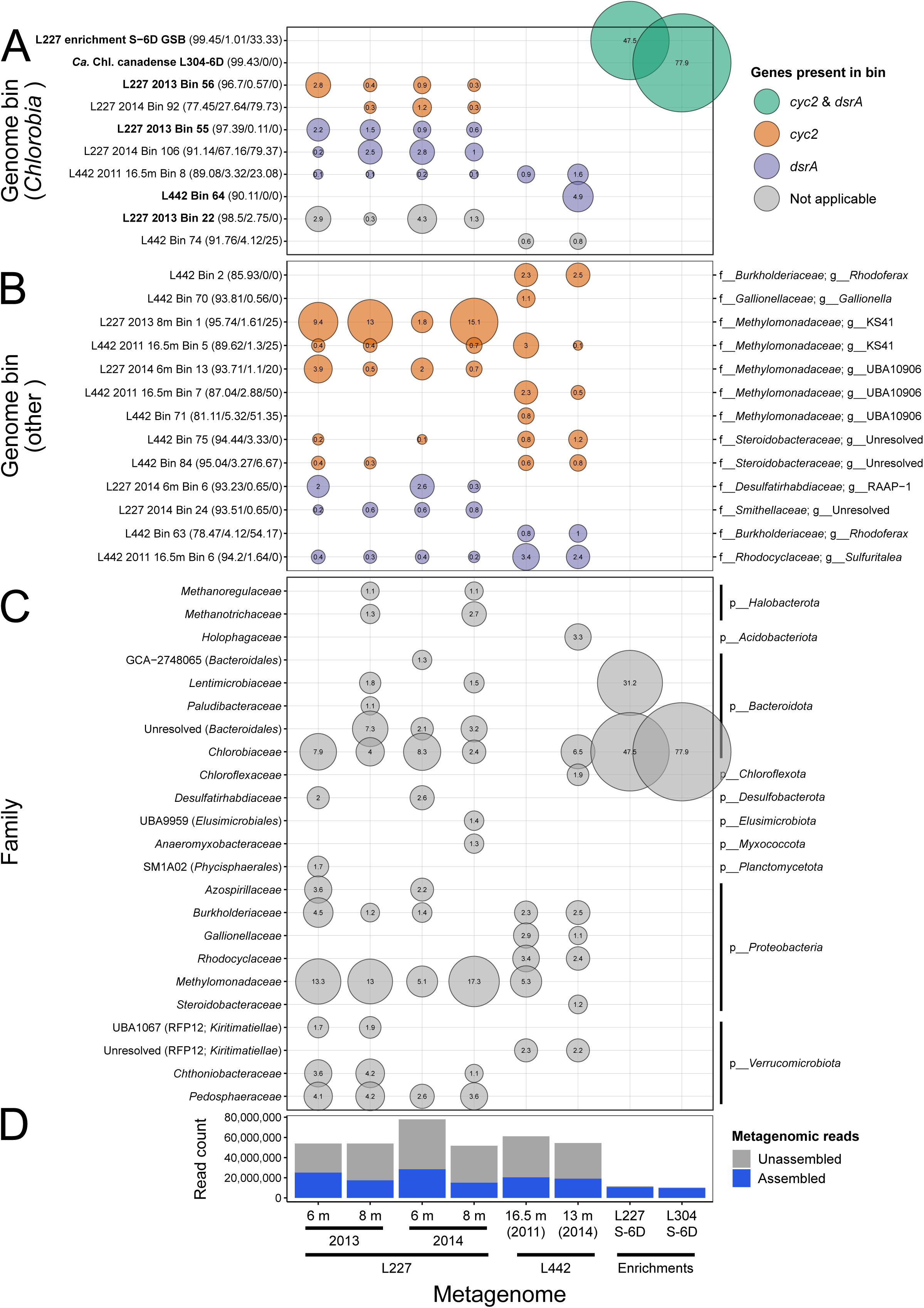
Bubble plot showing predicted relative abundances of recovered genome bins in the lake and enrichment culture environments. The size of each bubble corresponds to the relative abundance of a genome bin or taxon within metagenome data based on mapping of assembled reads. Each bubble is also labelled with its percent relative abundance for clarity. **A**, Bins of *Chlorobia* coloured by their metabolic potential based on functional gene markers described in this study. Bolded names correspond to the higher-quality bins described in Table 1 and Fig. 2. In parentheses beside each name are bin quality statistics reported by CheckM – the predicted % completion, % contamination, and % strain heterogeneity, respectively. The displayed bins all classify to the *Chlorobium* genus based on GTDB taxonomy. **B**, Non-*Chlorobia* bins (≥ 0.01% relative abundance) found to have the same functional gene markers (*cyc2* or *dsrA*) in their genome sequences. Quality statistics are reported as in panel **A**. On the right side of the panel, the GTDB family and genus classifications of each bin are shown. **C**, Family-level taxonomic composition of the metagenomes based on GTDB classifications of genome bins with ≥ 1% relative abundance. The phylum of each family is displayed on the right side of the panel. For unresolved or non-Latin family names, the order (and class, if needed) is displayed in parentheses beside the family name for clarity. **D**, Assembly statistics for the metagenome data. The total number of quality-trimmed metagenome reads is represented by the total height of each bar. Reads that mapped to the filtered sequence assembly (i.e., excluding short contigs, see Methods) are highlighted in blue and are considered “assembled” reads. Assembled read totals were used to determine relative abundances of genome bins in panels A-C. Supplementary Figure 4 shows relative abundances based on predictions from unassembled metagenome reads for comparison.

Several non-*Chlorobia* genome bins were also found to contain the *cyc2* or *dsrA* gene through MetAnnotate-based HMM searches (Fig. 3B). Genome bins containing potential *cyc2* homologues grouped into the *Rhodoferax* or *Gallionella* genera known to include iron-cycling bacteria [18, 80] or into families not associated with iron oxidation (i.e., *Methylomonadaceae* and *Steroidobacteraceae*). The genome bins classified to the *Methylomonadaceae* family were at high relative abundance in the lake metagenomes, suggesting an abundance potentially as high as ∼17% in Lake 227 (Fig. 3C). All *Methylomonadaceae*-affiliated bins, except for L442 2011 16.5m Bin 5, encoded particulate methane monooxygenase (*pmoA*) genes. Genome bins containing *dsrA* included known chemolithotrophic sulfide-oxidizing bacteria of the *Sulfuritalea* genus, heterotrophic sulfate-reducing bacteria of the *Desulfatirhabdiaceae* family, a bin classified to the *Smithellaceae* family of the *Syntrophales* order, and a bin classified to the *Rhodoferax* genus [81, 82]. These taxa, with the exception of *Rhodoferax* and *Smithellaceae*, also appeared in unassembled read-based marker gene assessments of the metagenomes (Supplementary Figure 5) and could play a role in the broader iron and sulfur cycles of the lakes.

*Chlorobia* appeared to represent the dominant phototrophs in the lake anoxic zone samples. In total, genome bins classified to the *Chlorobia* represented as high as 8.3% of lake microbial communities based on read mapping (Fig. 3C). Predicted relative abundances of populations of *Chlorobia* were 1.2-1.8 times higher when unassembled reads were assessed directly (Supplementary Figure 5), with a predicted maximum relative abundance of 12.3%, showing that high relative abundance is not a result of assembly bias (Fig. 3D). The only other potential chlorophototroph detected in the dataset at >1 % relative abundance was a single genome bin classified to the *Chloroflexaceae* family (L442 Bin 82; 92.9/1.2% completeness/contamination), which made up 1.9% of the L442 microbial community in 2014 at 15 m depth (Fig. 3C) and contained the *bchL* and *pufM* photosynthesis genes (data not shown). This genome bin was undetected in all other samples.

### Assessment of ferrous iron oxidation potential of “Ca. Chl. canadense”

Cultures of *Chl. ferrooxidans* showed the expected behaviour when incubated in ferrous iron containing medium. After a short lag phase, both *Chl. ferrooxidans* cultures containing 100 μM FeCl_2_ (no EDTA) and exposed to light began oxidizing ferrous iron, and the cultures had nearly completely oxidized all ferrous iron within two days of the initial setup (Fig. 4). Similar bottles incubated in the dark showed consistent iron oxidation rates of ∼5 μM d^−1^ that corresponded to rates observed in uninoculated media controls; this iron oxidation was inferred to be abiotic, for example due to small amounts of oxygen contamination during sampling. No ferrous iron oxidation outside of this baseline effect was observed in the *Chl. ferrooxidans* cultures supplemented with 120 μM EDTA, in contrast to the stimulatory effect of EDTA reported for *Rhodopseudomonas palustris* TIE-1 [74], which uses a different gene system (*pioAB*) for extracellular electron transfer [19]. The EDTA-amended medium consistently appeared to have ∼50% ferrous iron oxidation at the beginning of the experiment, based on the ferrozine assay, but this apparent oxidation has been observed previously with EDTA and could potentially be due to ineffective binding of ferrozine to EDTA-chelated iron [76].

**Figure 4.**
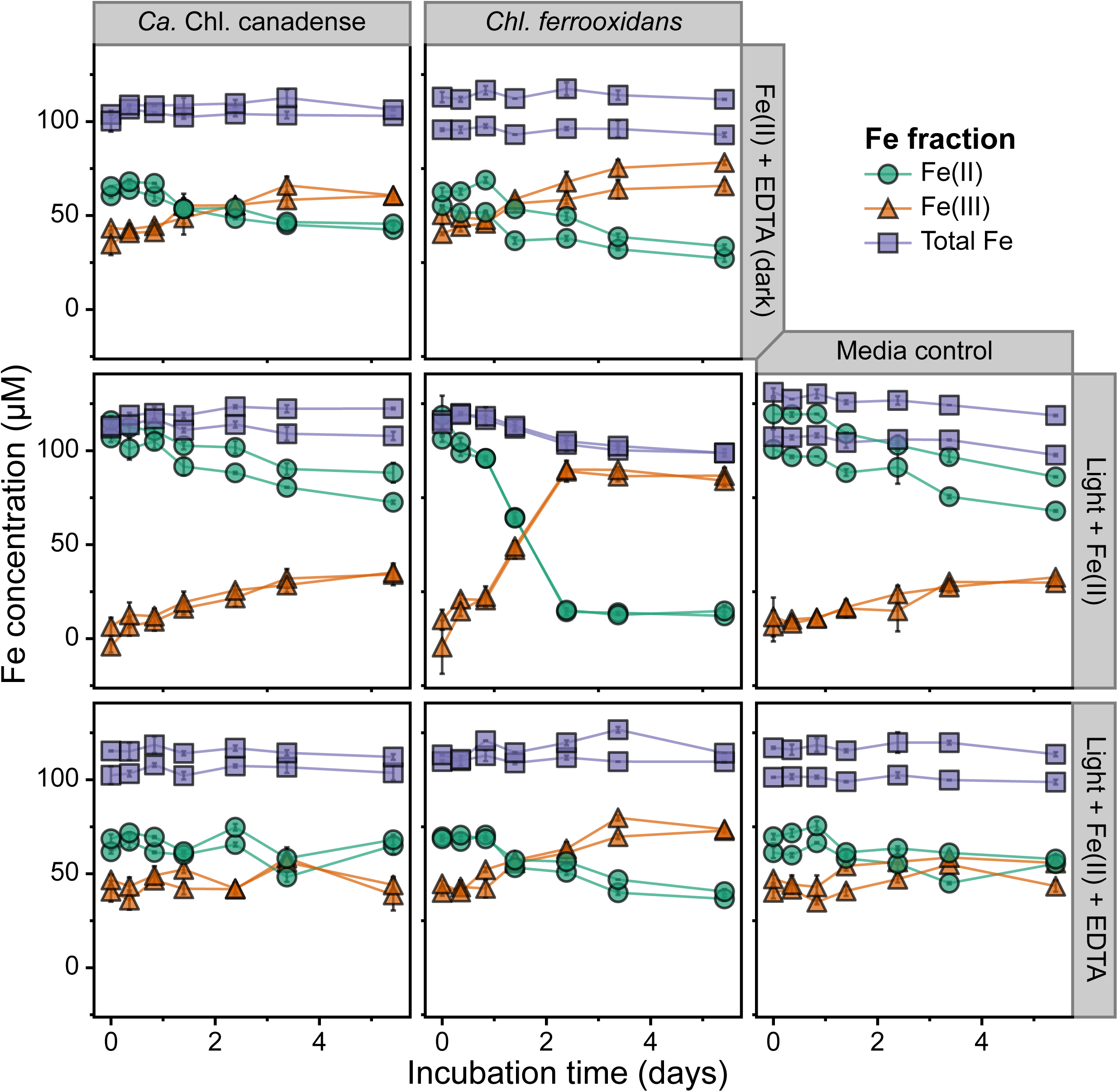
Iron oxidation activity of two *Chlorobia* cultures under different growth conditions over time. The concentrations of ferrous iron, total iron, and ferric iron (determined by the subtraction of ferrous iron from total iron) are shown over a 10 day incubation period for “*Ca.* Chl. canadense” (enriched in this study) and *Chl. ferrooxidans*. The activity of the cultures in iron-amended media is shown when incubated in the dark with EDTA, in the light, and in the light with EDTA. Each panel depicts the results of biological duplicates, with error bars representing the standard deviation of technical duplicates in the ferrozine assay. The same results are shown for the full 21-day incubation period in Supplementary Figure 6.

In contrast to *Chl. ferrooxidans*, “*Ca.* Chl. canadense” showed no photoferrotrophic ferrous iron oxidation activity over the course of the experiment (Fig. 4; Supplementary Figure 6), and the iron oxidation profiles of the cultures closely matched those of the uninoculated media control bottles. A reference bottle of Hegler freshwater medium prepared in the same batch as those in the main experiment but amended with 600 μM buffered sulfide feeding solution, in place of ferrous iron, received the same inoculum of “*Ca.* Chl. canadense”. Complete oxidation of sulfide in the bottle was observed within five days of inoculation and incubation under light (not shown), indicating that the media and inoculum were not the cause of the lack of observed photoferrotrophic activity.

## Discussion

Our study offers genome-resolved functional gene evidence and enrichment cultivation data that provides a first glimpse into the ecology of photoferrotrophy in seasonally anoxic Boreal Shield lakes. These results represent the first application of genome-resolved metagenomics to an Archaean ocean analogue system and the first use of this approach to delineate potential metabolic modes of coexisting, closely related photoautotrophs. We suggest the genomic potential for photoferrotrophy in Lake 227 based on detection of the *cyc2* gene in multiple genome bins of *Chlorobia*, and we argue for complex phototrophic ecology in Boreal Shield lakes based on metabolic diversity of photoautotrophs within and between lake systems. Overall, our findings provide genomic context to previous 16S rRNA gene data supporting the existence of photoferrotrophy in Boreal Shield lakes [37], adding nuance to the interpretation of these data while expanding sparse knowledge of the diversity of *cyc2* within the class *Chlorobia*. Our research also reinforces the need for additional genetic work to understand the function and regulation of *cyc2* in photoferrotrophy.

*Chlorobia*-affliated *cyc2* genes were detected in high relative abundance genome bins in Lake 227 metagenomic data. The dominant *cyc2*-containing genome bin of *Chlorobia* in Lake 227 metagenomic data, L227 2013 Bin 56, was highly complete and represented as much as 2.8% of the upper anoxic zone microbial community based on read recruitment (Fig. 3A). Although populations of *Chlorobia* are commonly detected at high relative abundances in the anoxic zones of boreal lakes, these populations are typically assumed to oxidize sulfide (e.g., [83]). Because L227 2013 Bin 56 encoded no detectable genes for sulfide or thiosulfate oxidation (Fig. 2), the corresponding population might rely solely on *cyc2* to supply electrons (e.g., from ferrous iron) needed for photosynthesis. Although the population represented by this genome bin might obtain energy chemotrophically via hydrogen oxidation (Fig. 2), as is possible for many members of the *Chlorobia* class [25], the higher relative abundance of *Chlorobia* in the upper compared to lower anoxic zone of Lake 227 (based on previous 16S rRNA gene sequencing data [37]) suggests that light energy is being used in metabolism. The lack of detected sulfur oxidation genes in L227 2013 Bin 56 might be due to incomplete binning, given that the bin was found to encode only partial gene pathways for sulfate assimilation and the reverse TCA cycle (Fig. 2; Supplementary Figure 3), and additional work would be needed to confirm whether *cyc2* is actively expressed. However, our detection of *cyc2* affiliated with this high relative abundance genome bin of *Chlorobia* in Lake 227 metagenomes provides important first evidence that the gene may bear ecological relevance for this Boreal Shield lake.

Our results reveal the coexistence of populations of *Chlorobia* with varying metabolic potential in Boreal Shield lakes. Genome bins of *Chlorobia* encoding *cyc2* co-occurred consistently with bins of *Chlorobia* encoding sulfide oxidation genes (and lacking *cyc2*) at similar relative abundances across all four analyzed Lake 227 metagenomes (Fig. 3A). Such coexistence aligns well with observations of co-occurring sulfide-oxidizing phototrophs and photoferrotrophs in Lake La Cruz [34] and agrees with existing evidence that phototrophic sulfide oxidation can occur in ferruginous systems [33]. Observations from enrichment culturing on ferrous iron-containing media suggest that sulfide oxidation among Boreal Shield lake *Chlorobia* is possible even when sulfide is at trace levels, generated directly from sulfate-reducing bacteria (Supplementary Figure 2). Such “cryptic sulfur cycling” has been reported in various other aquatic systems and is likely an ecologically common phenomenon beyond Boreal Shield lakes [84] and common among phototrophic processes even within low-sulfate environments. Cryptic iron cycling, which has been observed in other ferruginous lakes [36], might also be possible in Boreal Shield lakes but was not observed directly in this study.

The lack of detectable *Chlorobia*-affiliated *cyc2* genes in Lake 442 metagenomes has important implications for the ecology of *cyc2*-encoding *Chlorobia*. Although genome bins of *Chlorobia* were estimated to occur at high relative abundance in Lake 442 based on metagenome data (i.e., 6.5% at 15 m depth in 2014), genome bins only encoded sulfide oxidation genes, not *cyc2* (Fig. 3A). Lake 442 has similar underlying geology to Lake 227 and is nearby (∼13.5 km distance). This suggests that the favourability of *cyc2*-encoding *Chlorobia* may be strongly influenced by physicochemical drivers within the natural range possible among Boreal Shield lakes. Generally, levels of sulfate in the water column are higher in Lake 442 than Lake 227 (i.e., ∼15 µM in the anoxic zone compared to ∼7.5 µM [37]), and total dissolved iron concentrations in Lake 442 tend to be lower at the depths where populations of *Chlorobia* are present (i.e., ∼5-15 µM compared to ∼10-100 µM in Lake 227). Lake 304, from which the *cyc2-*encoding “*Ca.* Chl. canadense” was enriched, has sulfate and total dissolved iron concentrations in its upper anoxic zone (∼10 uM and ∼40-200 uM, respectively; unpublished data) that are closer to Lake 227 values than Lake 442 values. It is possible that these factors, or other factors such as hydrogen and bicarbonate concentrations (observed to impact photoferrotrophy rates in *Rps. palustris* TIE-1 [85]), lake bathymetry, anoxia timing, pH, light quality, or dissolved organic carbon levels, could shift the geochemical balance in favour of *Chlorobia* encoding genes for phototrophic sulfide oxidation rather than *cyc2* in Lake 442. Although Lake 304 was historically fertilized until 1976, it is more likely, given Lake 304’s rapid recovery after fertilization ceased and ∼2.7 year water residence time (see Methods), that other similarities between Lakes 227 and 304, such as lake bathymetry [86] or high sediment fluxes of phosphorous observed in Lake 304 even before artificial nutrient addition began [87], allow for the favourability of *cyc2-*encoding *Chlorobia* in these lakes more than legacy fertilization effects. Understanding key factors governing the favourability of *cyc2-*associated metabolism among a broader suite of Boreal Shield lakes could prove highly valuable for predicting photoferrotrophic activity in both modern and Archaean ocean settings.

The lack of *Chlorobia*-affiliated *cyc2* genes in Lake 442 metagenomes also has implications for the interpretation of iron isotope data. The strong iron isotope fractionation observed between the oxic and anoxic zone boundaries of Lakes 227 and 442 previously by Schiff and colleagues [37] appears to occur regardless of whether *Chlorobia*-affiliated *cyc2* genes are detectable, implying that this signature cannot be used as a sole indicator of photoferrotrophic metabolism. It is possible that microaerophilic iron oxidation could contribute to the observed iron isotope fractionation in Lake 442 given the detection of a *cyc2*-encoding genome bin within the *Gallionelles* genus in Lake 442 metagenomic data (Fig. 3B). However, other yet unknown iron cycling processes might also be at work. More nuanced physico-chemical measurements or knowledge of additional biological processes are likely needed to robustly predict the potential for photoferrotrophy in modern and potentially ancient ferruginous environments.

We were unable to induce photoferrotrophy in “*Ca.* Chl. canadense” (Fig. 4), which was enriched in the present study from Lake 304. This new species, along with the Lake 227 enrichment that was lost in early subculturing, encoded both the *cyc2* gene and the sulfide oxidation gene pathway, suggesting an ability to switch between phototrophic sulfide oxidation and photoferrotrophy, as observed for *Chl.* sp. N1. A lack of observed photoferrotrophic growth in “*Ca.* Chl. canadense” might be a result of several factors. We noted that the c5 family cytochrome encoded by “*Ca.* Chl. canadense” directly upstream of *cyc2* contains an additional cysteine residue in the heme-binding motif in its primary sequence (i.e., CCXCH rather than the typical CXXCH), unlike the primary sequences of the c5 family cytochromes in reference cultures or environmental genome bins (Supplementary File 6). This additional cysteine might not completely compromise the function of the protein but could have deleterious consequences for iron oxidation [88]. If this gene neighbour to *cyc2* is involved in the same metabolic pathway as *cyc2*, it could imply that *cyc2* in “*Ca.* Chl. canadense” has an impaired or alternative function. Alternative functions could include electron exchange with humic substances as hypothesized by He and colleagues [89] or even manganese oxidation as reported recently for *Chlorobia* in mixed culture [90], although direct links between Cyc2 and oxidation of alternate substrates to ferrous iron have not been demonstrated experimentally. Cyc2 might also allow for direct electron uptake from solid-phase conductive substances, as has been observed in the photoferrotroph *Rps. palustris* TIE-1 [91]. Some of these alternative potential functions of Cyc2 might be preserved or even enhanced even if iron oxidation potential is impaired. Given that strong ecological controls appear to exist on *cyc2-*encoding *Chlorobia* (see above), it is also possible that the *cyc2* gene product in “*Ca.* Chl. canadense” is indeed involved in iron oxidation but requires carefully tuned conditions to switch metabolic modes. This option would align with current research that implicates *cyc2* in iron oxidation among neutrophilic iron-oxidizing bacteria (e.g., [20]). It would also align with preliminary reports that the *cyc2-*encoding but sulfide-oxidizing *Chl. luteolum* is capable of changing to photoferrotrophic growth (Kate Thompson and Sean Crowe, personal communication), despite previous difficulties inducing this behaviour [23]. The “*Ca*. Chl. canadense” culture provides a unique opportunity to probe the regulation and function of *cyc2* within the class *Chlorobia* in future work.

The addition of the genome bins of *Chlorobia* from this study to existing reference genomes doubles the known diversity of *Chlorobia*-associated *cyc2* genes and allows for further examination of the evolutionary history of *cyc2* in the class *Chlorobia*. Rather than forming a single monophyletic grouping, the presence of *cyc2* in genomes of *Chlorobia* appears polyphyletic based on the ribosomal protein phylogeny (Fig. 2), interspersed with genomes encoding various forms of sulfur oxidation. The thiosulfate oxidation (*sox*) gene system among *Chlorobia* is known to not follow an inheritance pattern matching taxonomic marker genes, and lateral gene transfer within the *Chlorobia* may have been an important factor in its acquisition [25, 92]. However, the *cyc2* predicted primary sequence phylogeny among *Chlorobia* (Fig. 1C) is somewhat congruent to the *Chlorobia* ribosomal protein phylogeny, suggesting that loss from a single common ancestor, rather than extensive lateral gene transfer within the class, may account for the majority of how *cyc2* genes are distributed within the *Chlorobia* class (Supplementary Figure 4). Such a scenario would imply that *cyc2* was acquired by a common ancestor of modern *Chlorobia* but multiple clades of modern *Chlorobia* lost the *cyc2* gene over evolutionary time. Recovery of additional *cyc2*-containing *Chlorobia* genomes is required to confirm this hypothesis, but it is an intriguing example of the types of evolutionary analyses possible as *cyc2* among the *Chlorobia* is further studied at the genomic level.

Our research provides insight into the metabolic diversity and ecology of anoxygenic photoautotrophs in seasonally anoxic Boreal Shield lakes. We show that high relative abundance genome bins of *Chlorobia* in lake metagenomes can have varying functional gene content with respect to their photosynthetic electron donor, including some genome bins from Lake 227 samples that encoded the *cyc2* gene implicated in photoferrotrophy. The absence of *Chlorobia-* affiliated *cyc2* genes in the neighbouring Lake 442 suggests that ecological or geochemical controls strongly influence the favourability of *cyc2*-based metabolism in Boreal Shield lakes, with potential impacts on whole microbial community function in the anoxic zone. Although we were unable to induce photoferrotrophy in a laboratory setting, we enriched the novel *cyc2-* encoding species “*Ca.* Chl. canadense”, which could be used in future research exploring the function and regulation of *cyc2* as genetic studies continue to progress in linking the *cyc2* gene product to its cellular role [93]. Probing the metabolic diversity of anoxygenic phototrophs in Boreal Shield lakes could lead to novel inferences about photosynthesis in early Earth oceans, including how environmental variables in the range possible for Archaean oceans could modulate the dominant mode of photosynthesis. Altogether, our findings serve as an important basis for future work probing the rates, gene expression, biogeography, and enrichment culture activity of *cyc2*-encoding *Chlorobia* in Boreal Shield lakes. Although further work is needed to demonstrate the role of the *cyc2* gene product in photoferrotrophy and the distribution of *cyc2* across a broader suite of Boreal Shield lakes, our data indicate that metabolic diversity among anoxygenic photoautotrophs may be an overlooked yet ecologically important phenomenon in Boreal Shield lakes globally. Knowledge of such metabolic diversity impacts our understanding of modern ecosystems and predictions about the evolution of early life on Earth.

## Data availability

Raw metagenome reads for the freshwater lake and enrichment culture metagenomes are available in the NCBI sequence read archive (SRA) under BioProjects PRJNA518727 and PRJNA534305, respectively. Six curated genome bins of *Chlorobia* are available under the same BioProjects, along with all assembled contigs from freshwater lake metagenomes. The complete set of uncurated genome bins were uploaded to a Zenodo repository at <u>doi:10.5281/zenodo.2720705.</u> Code for downloading these data and performing the analyses presented in this paper is available at https://github.com/jmtsuji/Chlorobia-cyc2-genomics (<u>doi:10.5281/zenodo.3228523</u>).

## Acknowledgements

We thank R. Henderson and staff at the IISD-ELA for assistance in field sampling and interpretation of the dynamics of ELA lakes. In addition, we thank S. A. Crowe for valuable feedback on the manuscript, K. J. Thompson for helpful discussion related to the work, L. H. Bergstrand for assistance with bioinformatics, and K. E. Engel, R. C. Beaver, and N. A. Shaw for assistance with laboratory tasks. JMT thanks M. S. M. Jetten, C. U. Welte, and staff at the Soehngen Institute of Anaerobic Microbiology (SIAM), for providing helpful training for anaerobic microbiology methods. This research was supported by Discovery and Strategic Projects Grants from the National Sciences and Engineering Research Council of Canada (NSERC).

## Author contributions

SLS, JJV, and LM were involved in the overall lake sampling project that led to this work and provided input and assistance with sample collection and data interpretation. JMT and JDN designed the experimental work in this study. JMT, NT, and MT performed enrichment cultivation with support from SH. JMT analyzed the sequence data and drafted the manuscript with the edits and feedback of all authors.

## Supplementary information

### Supplementary Methods

Supplementary Figure 1 – Average nucleotide identity between recovered genome bins and reference genomes of *Chlorobia*. The same ribosomal protein tree as in Figure 2 is shown to the left of the heatmap.

Supplementary Figure 2 – Photographs of enrichment cultures of *Chlorobia* in ferrous iron- and sulfide-containing media.

Supplementary Figure 3 – Genomic potential for photosynthesis and carbon fixation in recovered genome bins of *Chlorobia* compared to reference strains. The figure layout and phylogenetic tree are identical to Figure 2. All query sequences for reciprocal BLASTP were derived from the genome of *Chl. tepidum* TLS except for BciB, which was derived from the genome of *Chl. ferrooxidans* KoFox.

Supplementary Figure 4 – Tanglegram comparing the concatenated ribosomal protein phylogeny (Fig. 2) and Cyc2 phylogeny (Fig. 1C) among *Chlorobia*. The two tips whose placements differ between the two phylogenies are highlighted in blue or green.

Supplementary Figure 5 – Bubble plot showing taxonomic and functional profiling of unassembled metagenome data, in the same general layout as Figure 3. HMM hit counts were normalized to the total hits of *rpoB*, a single-copy taxonomic marker gene, within each sample, such that HMM hits are expressed as a percentage of *rpoB*. All families (NCBI taxonomy) with > 1% of normalized hits to the HMMs are shown. Although not shown, normalized *Chlorobiaceae*-associated *cyc2* hits for Lake 227 at 8 m were 0.9% for both 2013 and 2014.

Supplementary Figure 6 – Iron oxidation activity of cultures of *Chlorobia* over an extended incubation period of 21 days. The figure layout is identical to Figure 4.

Supplementary File 1 – ZIP file containing the YAML config files for the ATLAS metagenome assemblies performed for this publication.

Supplementary File 2 – ZIP file of two alternative profile Hidden Markov Models for the *cyc2* gene (general or *Chlorobia*-specific).

Supplementary File 3 – Excel file containing a summary of basic physical, sampling, and historic parameters for Lakes 227, 442, and 304.

Supplementary File 4 – Excel file containing accessions of reference genomes of *Chlorobia* and statistics from the reciprocal BLAST search of Fe/S gene pathways.

Supplementary File 5 – CSV file containing genome statistics for all dereplicated MAGs from this study.

Supplementary File 6 – ZIP file of amino acid multiple sequence alignments: alignment of *Chlorobia*-associated Cyc2 predicted primary sequences from this study with reference sequences (unmasked and masked), alignment of c5 family cytochromes presented in this study, and alignment of additional low-homology Cyc2 hits from metagenomes.

Supplementary File 7 – ZIP file of purity data for the “*Ca.* Chl. canadense” culture, including an epifluorescence microscopy image of the culture and an amplicon sequence variant (ASV) table summarizing the percent relative abundances of members of the enrichment culture.

## Table and figure legends

**Table 1.** Quality statistics for genome bins of *Chlorobia* recovered in this study. Completeness and contamination were calculated using CheckM (see Methods). Note that no ribosomal RNA genes were recovered for the bins. A summary of bin statistics for lower-quality genome bins is included in Supplementary File 5.

