## Supplementary Methods for "Genomic potential for photoferrotrophy in a seasonally anoxic Boreal Shield lake"

Running title: Photoferrotrophy in a Boreal Shield lake

**Supplementary methods**

*Co-assembly and binning*

Co-assembly and binning of lake metagenomes used a simple wrapper around the ATLAS pipeline, *co-assembly.sh*, which is available in the atlas-extensions GitHub repository at https://github.com/jmtsuji/atlas-extensions. Briefly, the wrapper combines QC processed reads from the original ATLAS run for samples of interest, re-runs ATLAS on the combined reads, maps QC processed reads from the original samples onto the co-assembly, and then uses the read mapping information to guide genome binning. Version 1.0.22-coassembly-r3 of co-assembly.sh was used, relying on identical settings to the original ATLAS run (see config file in Supplementary File 1), except that MEGAHIT was used for sequence assembly in place of metaSPAdes [1, 2], MetaBAT2 version 2.12.1 was used for genome binning [3], and, for the L227 coassembly, a contig length threshold of 2200 was used.

*Enrichment cultivation*

Enrichment cultures were maintained and purified in a variety of ways. Following initial enrichment, cultures were transferred with 1-10% inoculum into fresh media 2-4 times to continue to promote growth of the target phototroph. (Lake 227 enrichment S-6D was lost during these initial transfers.) Dilutions to extinction were then performed in liquid culture with dilution factors ranging from $10^{-2}$ to $10^{-7}$ to eliminate contaminating organisms. Later, deep agar dilution series was performed on the same cultures to further enrich the target organisms (see methods). Cultures were then transferred back to growth in liquid to yield higher cell biomass, with 5-10% inoculum typically being used in transfers between liquid cultures. In total, for "*Ca.* Chl.

canadense", 13 subcultures were performed from the initial lake water inoculum until the iron oxidation experiment (see methods) was performed.

*Phylogeny reconstruction from cyc2*

Prior to building the *cyc2* phylogeny, due to the poor sequence homology across much of the C-terminal end of the *cyc2* gene, phylogenetically uninformative residues in the alignment were masked using Gblocks, version 0.91b, with the flags "-t=p -b3=40 -b4=4 -b5=h", reducing the alignment size from 609 to 223 residues [4]. The phylogeny was then prepared from the masked sequence alignment via IQ-TREE, version 1.6.10 [5] as described in the main manuscript text.

*Comparison of ribosomal protein and cyc2 phylogenies*

To compare the ribosomal protein phylogeny and *cyc2* phylogeny for *Chlorobia* genomes containing *cyc2*, the amino acid sequence alignments used to construct the full phylogenies were subsetted to six relevant taxa and re-aligned. Maximum likelihood sequence phylogenies were constructed using IQ-TREE as described in the methods section for the full phylogenies. The tanglegram plot was visualized using Dendroscope version 3.5.10.

*Metagenome taxonomic and functional profiling*

Environmental relative abundances of microbial populations were estimated by read mapping to dereplicated genome bins. The QC processed metagenome reads from each sample were iteratively mapped to all dereplicated genome bins (> 75% completeness, < 25% contamination) using bbmap version 38.22 [6] to determine the proportion of read recruitment to each genome bin. To minimize read mapping from closely related strains, bbmap was run with the "*perfectmode*" flag so that only identical reads would map; all other settings were identical to those in the ATLAS config file in Supplementary File 1. Read recruitment was expressed in terms of the number of mapped metagenome reads to a genome bin divided by the total number of metagenome reads that mapped to assembled contigs. Overall, bin relative abundances are likely underestimated based on bbmap settings, which prevent SNPs from being detected, but are likely overestimated based on the calculation of read recruitment to assembled reads rather than total reads.

As a cross-comparison to the above genome bin-based method, pre-assembled metagenome reads were directly assessed for gene relative abundances. Open reading frames were predicted for QC processed metagenome reads using FragGeneScanPlusPlus commit 299cc18 [7]. Predicted open reading frames from the pre-assembled reads were then used with MetAnnotate development release version 0.9.2 [8] to scan for genes of interest using the HMM queries mentioned above and to classify the hits taxonomically. MetAnnotate performed taxonomic classification using the USEARCH method against the RefSeq database (release 80; March 2017). The default e-value cutoff of $10^{-3}$ was used for assigning taxonomy to hits. Relative abundances of phylotypes were calculated and visualized based on MetAnnotate results using *metannotate_barplots.R* version 1.1.0, available at https://github.com/jmtsuji/metannotate-analysis, with a HMM e-value cutoff of $10^{-10}$ to accommodate the shorter lengths of HMM hits.

*Assessment of ferrous iron oxidation potential of Chlorobia enrichments*

Generally, acidified samples for the ferrozine assay were stored at 4°C for less than two days before being assayed. As one exception, the acidified samples collected on day 14 (Supplementary Figure 6) were stored at 4°C for eight days before being assayed, yet no clear abnormalities of iron concentrations were observed for these samples compared to other samples in the time series.

To assess the purity of the culture, a positive control of "*Ca.* Chl. canadense" grown in a sulfide-containing medium, using the same inoculum as for the photoferrotrophy test, was examined using confocal laser scanning microscopy. A 5 mL aliquot of culture was pelleted, resuspended in 50 μL of supernatant, and visualized as a wet mount on a Zeiss LSM 700 confocal laser scanning microscope. To detect the autofluorescence of bacteriochlorophyll *c*/*d*/*e*-containing cells, the sample was excited using a 488 nm laser, and light emissions from 600-800 nm were measured. Transmitted light was also measured while imaging to provide information about the non-fluorescent portions of the slide.

In addition, for a previous subculture, genomic DNA was extracted from a pellet of cell biomass using the DNeasy UltraClean Microbial Kit, and 16S rRNA gene amplicon sequencing was performed, as described by Kennedy and colleagues [9], to identify the key contaminants in the culture. As an exception from the protocol by Kennedy and colleagues, the primers 515f and 926r (targeting the V4-V5 hypervariable region) were used for amplification [10], and PCR was performed in singlicate. During PCR, samples were incubated in the thermocycler at 95°C for 10 minutes, then incubated for 35 cycles of 95°C for 30 seconds, 50°C for 30 seconds, and 68°C for 1 minute, and finally incubated at 68°C for 7 minutes before being held at 12°C. Sequencing data was processed using QIIME2, version 2019.10.0 [11], within the AXIOME3 pipeline (https://github.com/neufeld/AXIOME3), development commit e35959d. Specifically, DADA2 was used to trim primer regions from raw sequencing data, merge paired end (2x250 base) reads, perform sequence denoising, and generate an amplicon sequencing variant (ASV) table [12]. Taxonomic classification of ASVs was performed using QIIME2's scikit learn-based taxonomy classifier against Silva release 132, trimmed to the V4-V5 region, as a reference database [13], and this information was overlaid on the ASV table. The Silva classifier was trained using QIIME2, version 2019.7.0, which relies on the same version of scikit learn as 2019.10.0.
