## Supplementary figures and images for "Genomic potential for photoferrotrophy in a seasonally anoxic Boreal Shield lake"

### Ca_Chl_canadense_S13.3a_wet_mount_autofluorescence.png

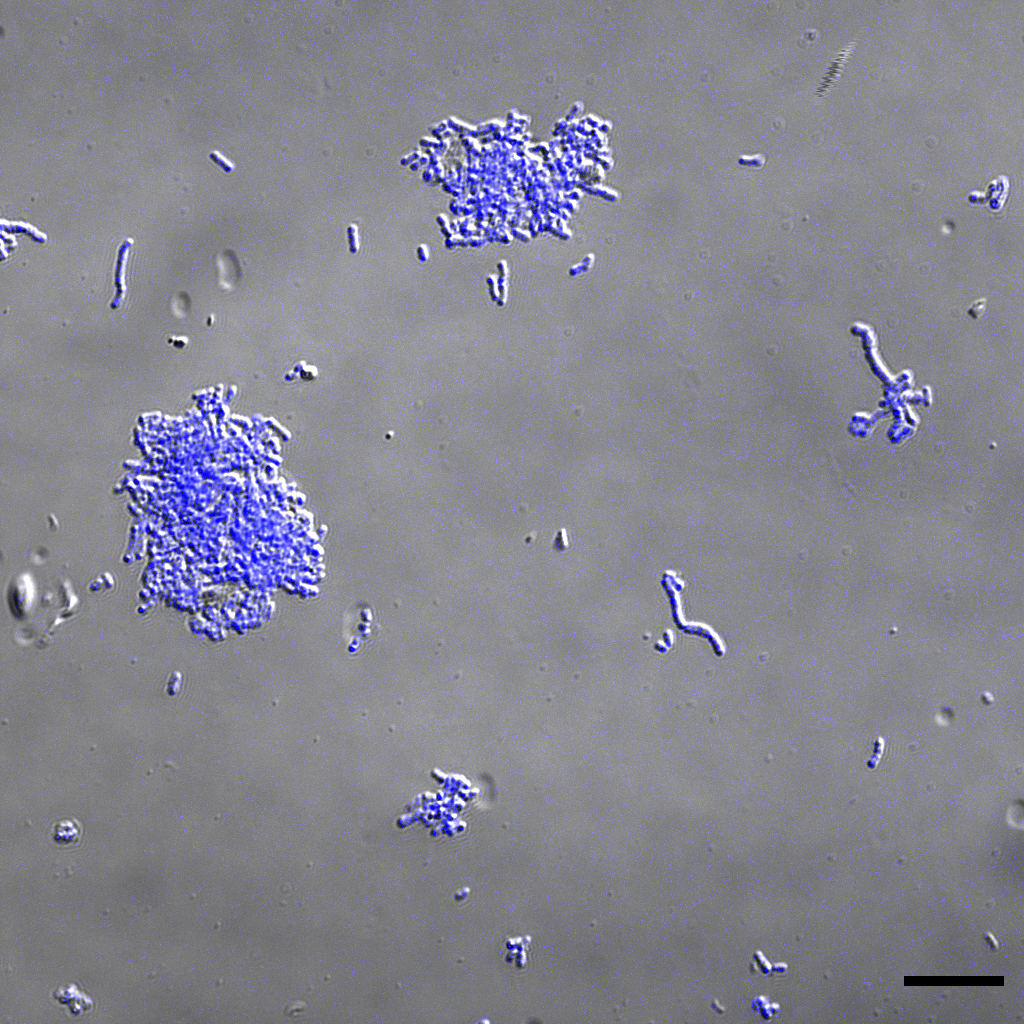

### Supplementary Figure 1

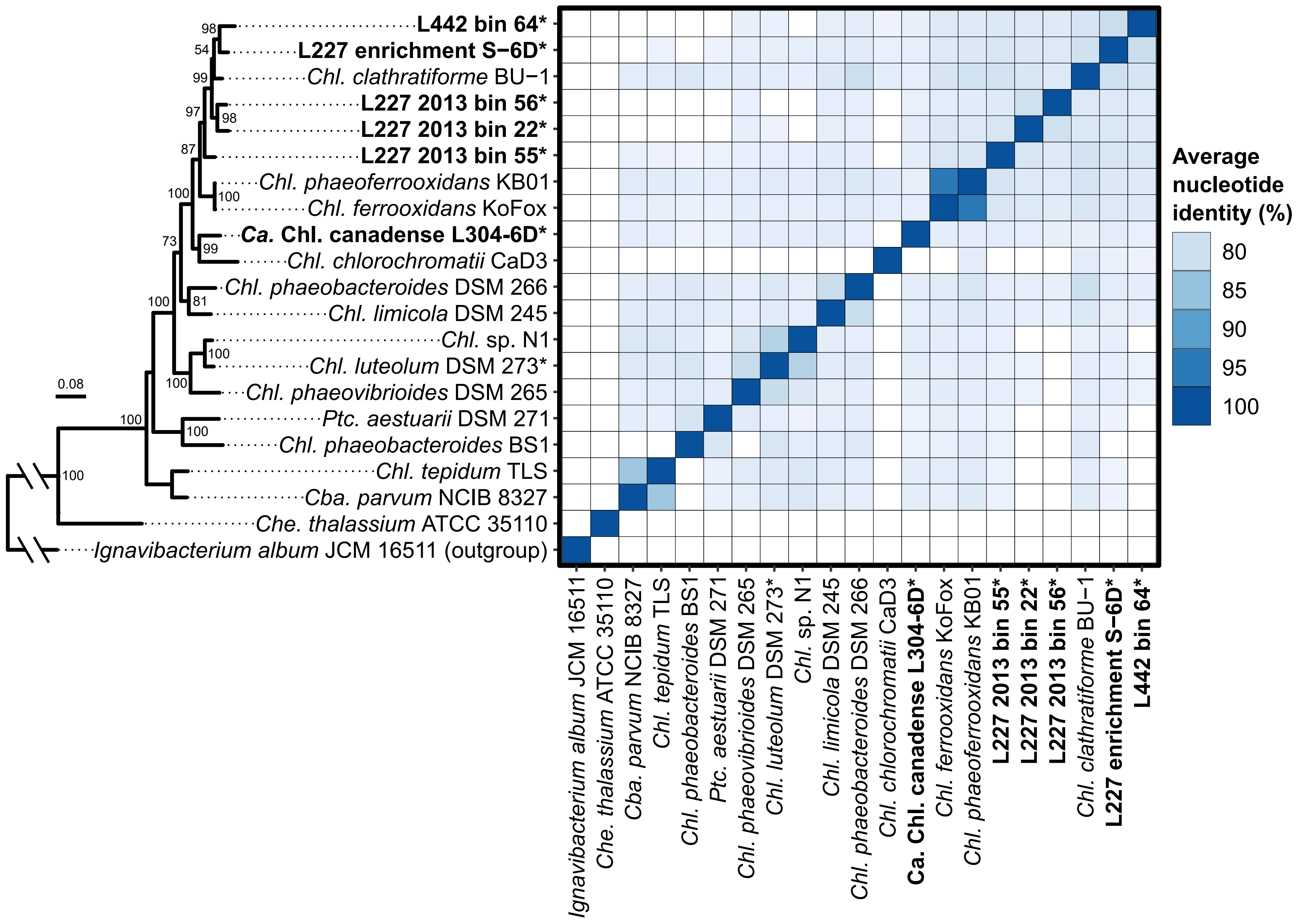

### Supplementary Figure 3

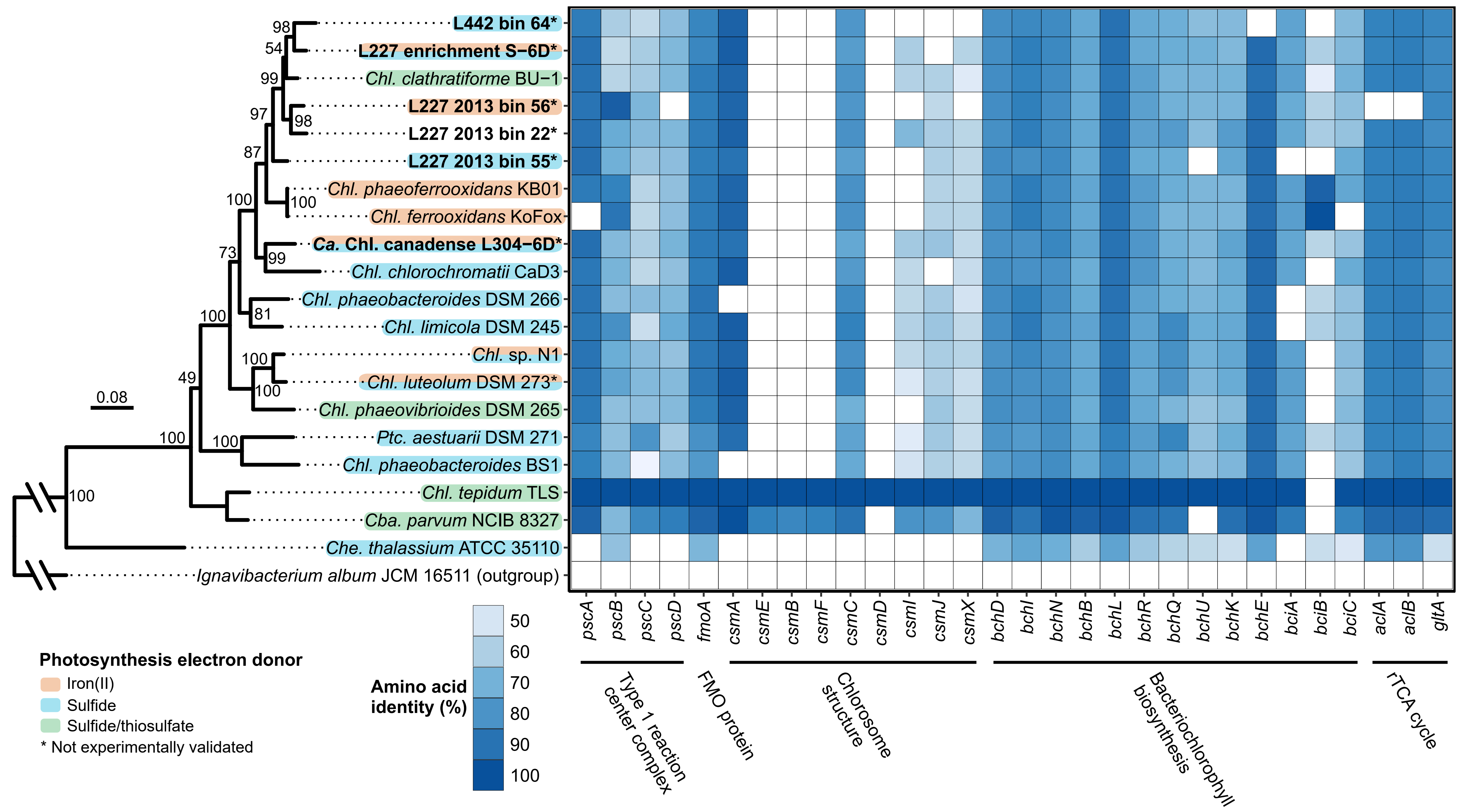

### Supplementary Figure 4

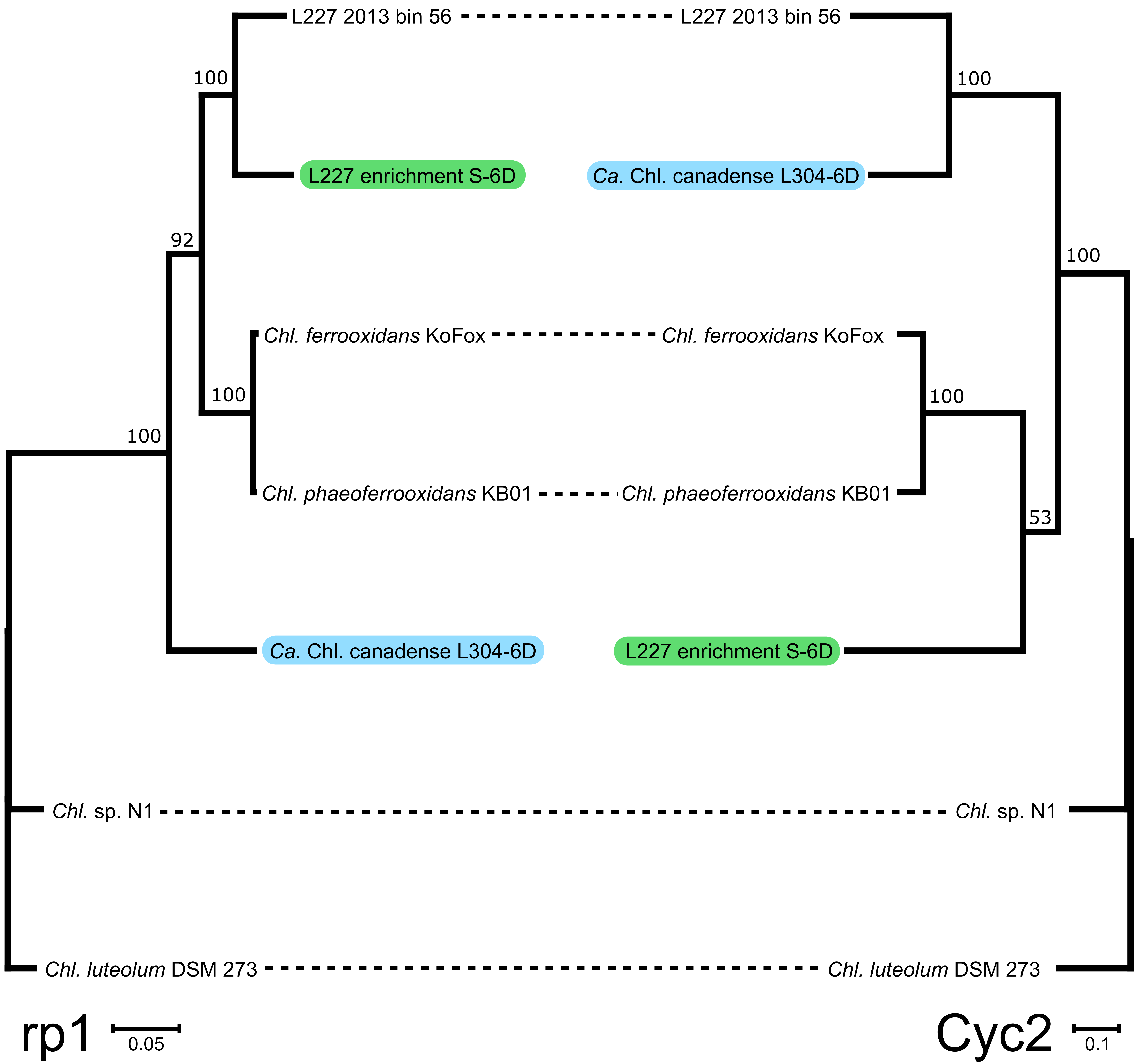

### Supplementary Figure 5

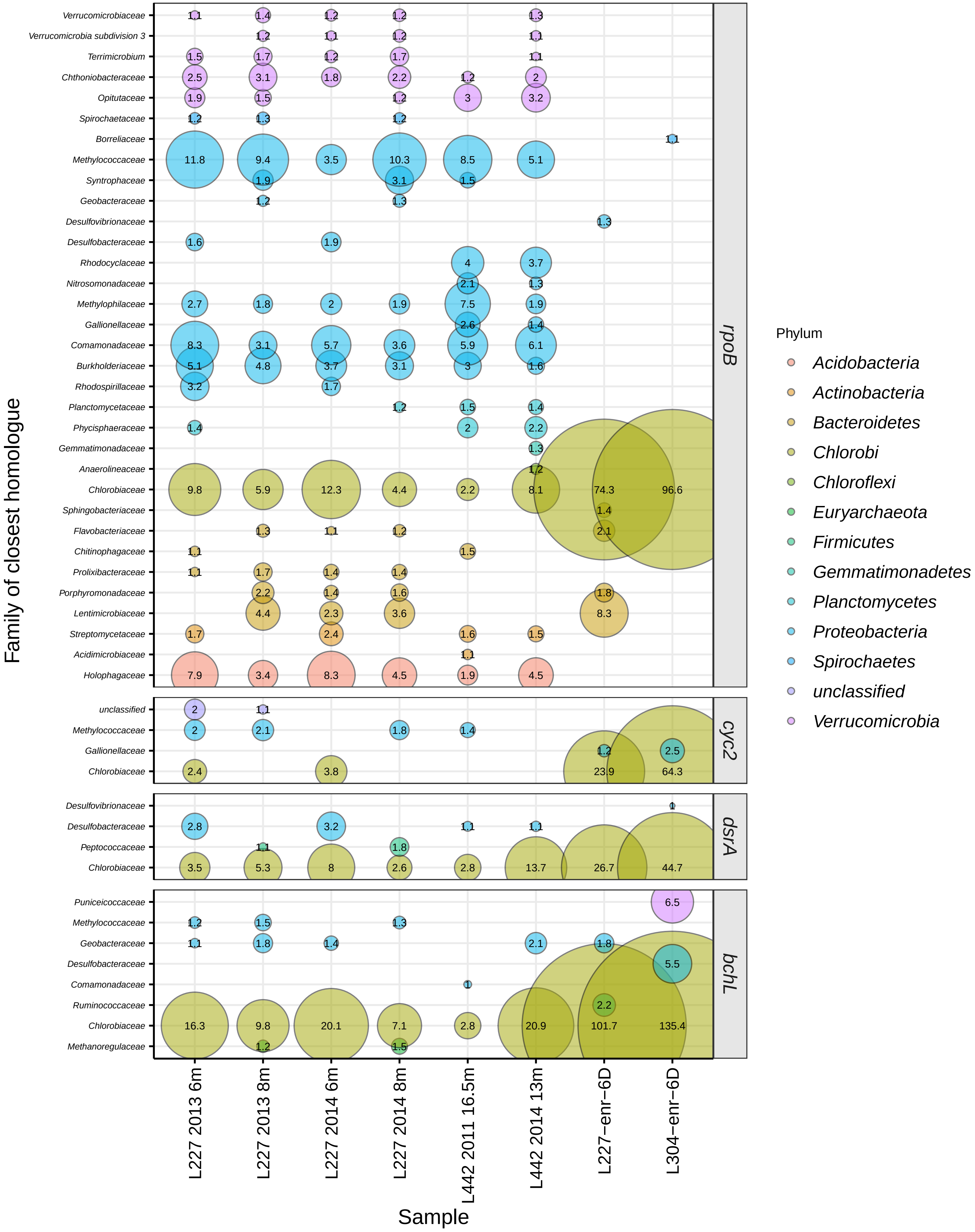

### Supplementary Figure 6

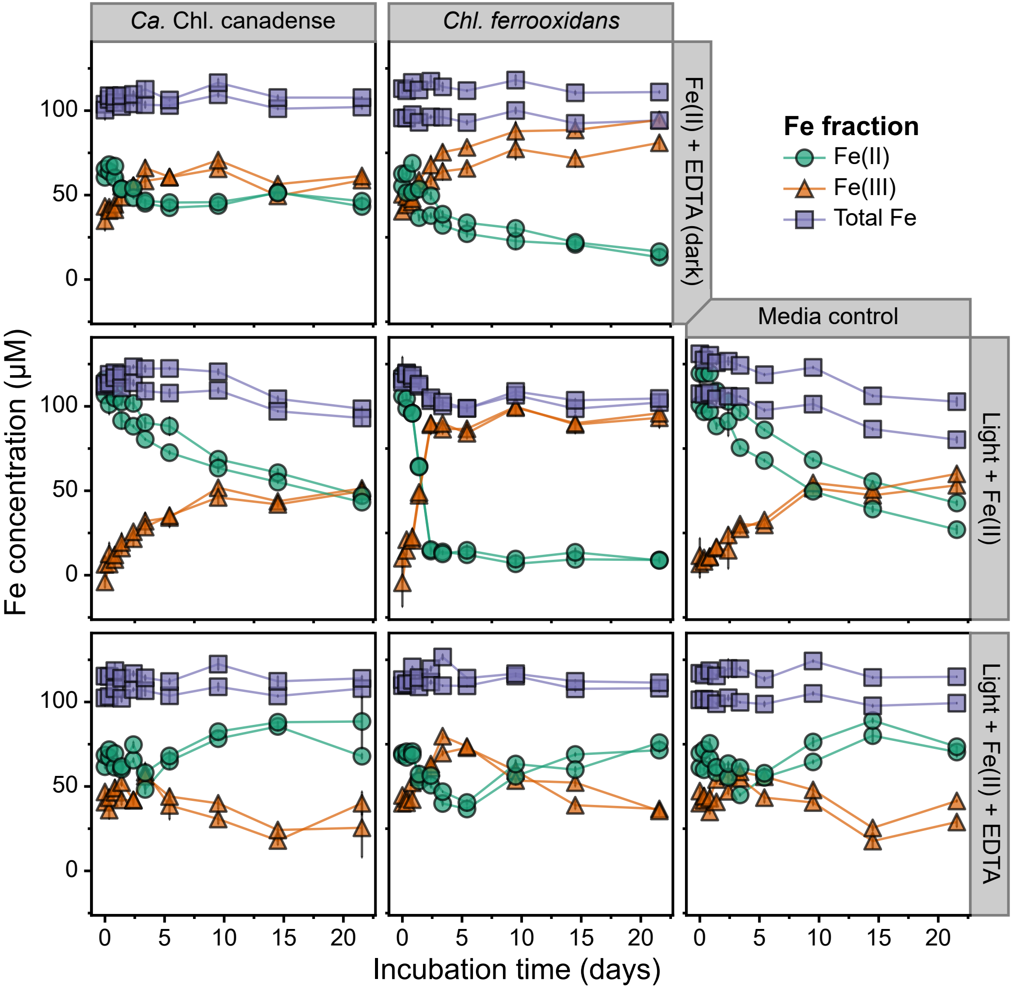
