## Supplementary Figure 2 for "Genomic potential for photoferrotrophy in a seasonally anoxic Boreal Shield lake"

### Photograph log of key enrichment cultures

**Media controls:**
Left: Pfennig's medium (S2-)

Right: Freshwater medium from Hegler and colleagues, 2008 ($Fe^{2+}$)

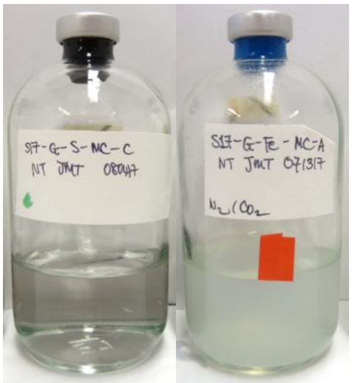

Reference culture of *Chl. ferrooxidans* KoFox grown on Hegler medium

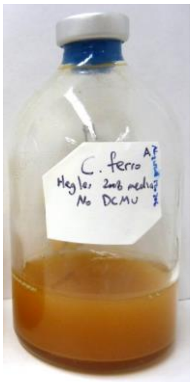

**L304 enrichment S-6D**
Inoculated Aug. 8th, 2017

Subcultured Nov. 23rd, 2017 (~3.5 months later)

Typical growth time of ~3 weeks

Cells ~0.8-1 x 1.2-1.5 µm in size

Named "*Ca.* Chl. canadense L304-6D"

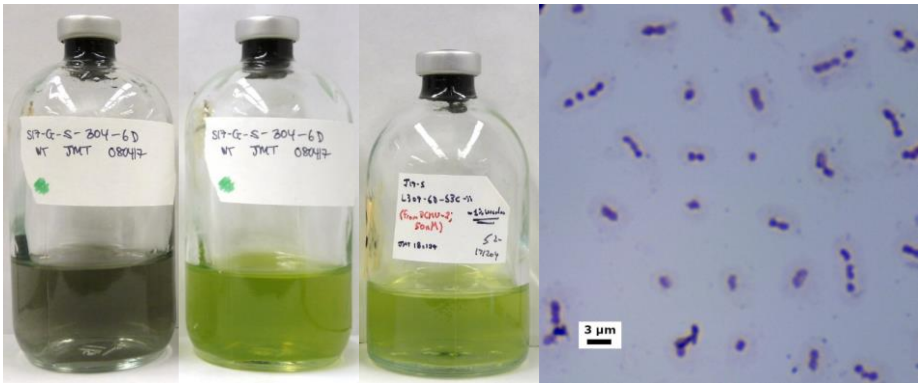


Aug. 24th, 2017 | Nov. 23rd, 2017 | Healthy subculture | Simple stain (dry mount); bright field microscopy

**L227 enrichment S-6D**
Dates of initial inoculation and subculture same as above

Typical growth time of ~4-8 weeks (slower growing)

Cells ~0.7-0.9 x 1-1.5 µm in size

**Strain was lost in May 2018**

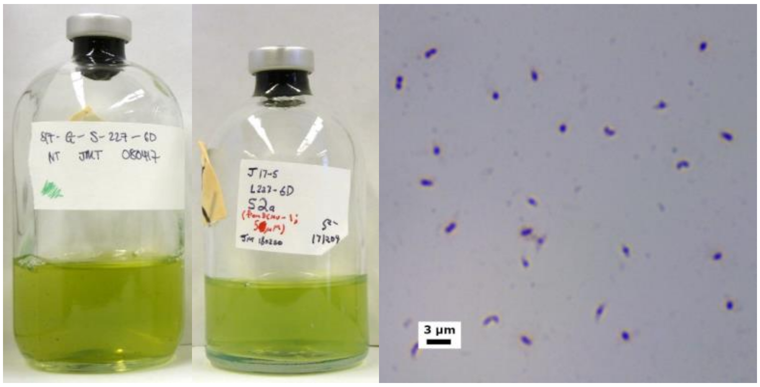


Nov. 23rd, 2017 | Healthy subculture | Simple stain (dry mount); bright field microscopy

**L227 enrichment Fe-6E**
Inoculated into Hegler medium ($Fe^{2+}$) on Aug. 8th, 2017

Developed greenish-black colouration

Eventually transitioned to sulfide-containing medium (Pfennig's), although growth was poor; *dsrA* sequence confirmed identical to L227-S-6D above

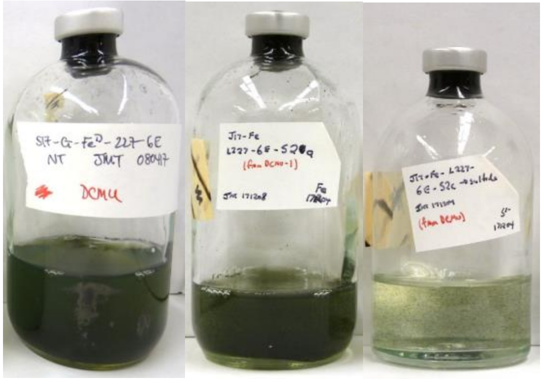

Oct. 19th, 2017 | Subculture, Hegler | Subculture, Pfennig
