## Supplementary File 1 for "Genomic potential for photoferrotrophy in a seasonally anoxic Boreal Shield lake": atlas_config_enrichments.yaml

```
samples:
  L227_S_6D:
    fastq:
    - /home/atlas/data/L227_S_6D_R1.fastq.gz
    - /home/atlas/data/L227_S_6D_R2.fastq.gz
    type: metagenome
  L304_S_6D:
    fastq:
    - /home/atlas/data/L304_S_6D_R1.fastq.gz
    - /home/atlas/data/L304_S_6D_R2.fastq.gz
    type: metagenome
tmpdir: /home/atlas/output/tmp
threads: 48
java_mem: 128
preprocess_adapters: /home/atlas/databases/adapters.fa
preprocess_adapter_min_k: 8
preprocess_minimum_base_quality: 10
preprocess_allowable_kmer_mismatches: 1
preprocess_reference_kmer_match_length: 27
preprocess_minimum_passing_read_length: 51
preprocess_minimum_base_frequency: 0.05
deduplicate: true
error_correction_overlapping_pairs: true
merge_pairs_before_assembly: true
contaminant_references:
  PhiX: /home/atlas/databases/phiX174_virus.fa
contaminant_max_indel: 20
contaminant_min_ratio: 0.65
contaminant_kmer_length: 13
contaminant_minimum_hits: 1
contaminant_ambiguous: best
duplicates_only_optical: false
duplicates_allow_substitutions: 2
normalization_kmer_length: 21
normalization_target_depth: 100
normalization_minimum_kmers: 15
merging_k: 62
merging_extend2: 50
merging_flags: ecct vstrict
assembler: spades
assembly_memory: 128
assembly_threads: 48
megahit_min_count: 2
megahit_k_min: 21
megahit_k_max: 121
megahit_k_step: 20
megahit_merge_level: 20,0.98
megahit_prune_level: 2
megahit_low_local_ratio: 0.2
minimum_contig_length: 1000
prefilter_minimum_contig_length: 500
spades_k: auto
minimum_average_coverage: 5
minimum_percent_covered_bases: 40
minimum_mapped_reads: 0
contig_trim_bp: 100
translation_table: 11
minimum_region_overlap: 1
feature_counts_allow_overlap: true
contig_count_multi_mapped_reads: false
contig_min_id: 0.76
contig_map_paired_only: true
contig_max_distance_between_pairs: 1000
maximum_counted_map_sites: 10
perform_genome_binning: true
maxbin_max_iteration: 50
maxbin_min_contig_length: 200
maxbin_prob_threshold: 0.9
refseq_namemap: /home/atlas/databases/refseq.db
refseq_tree: /home/atlas/databases/refseq.tree
diamond_db: /home/atlas/databases/refseq.dmnd
diamond_run_mode: fast
diamond_top_seqs: 2
diamond_e_value: 1.0e-06
diamond_min_identity: 50
diamond_query_coverage: 60
diamond_gap_open: 11
diamond_gap_extend: 1
diamond_block_size: 2
diamond_index_chunks: 4
summary_method: lca
aggregation_method: lca-majority
majority_threshold: 0.51
```
