## Supplementary File 2 for "Genomic potential for photoferrotrophy in a seasonally anoxic Boreal Shield lake": ABOUT_File_S2.txt

# ABOUT Supplementary File 2

This ZIP folder contains two files:

1. cyc2_all_references.hmm -- profile Hidden Markov Model built from an amino acid 
sequence alignment of the cyc2 genes recovered from Chlorobia in this study, the cyc2 
genes from the four reference Chlorobia, and the cyc2 genes summarized from reference 
genomes by He and colleagues, 2017 (see reference in main text)

2. cyc2_Chlorobia.hmm -- profile Hidden Markov Model built from an amino acid sequence 
alignment of the cyc2 genes recovered from Chlorobia in this study, the cyc2 genes from 
the four reference Chlorobia
