## Supplementary File 6 for "Genomic potential for photoferrotrophy in a seasonally anoxic Boreal Shield lake": ABOUT_File_S6.txt

# ABOUT Supplementary File 6

This ZIP folder contains four files:

1. cyc2_Fig1_unmasked.faa -- amino acid sequence alignment of the cyc2 genes recovered 
from Chlorobia in this study, the cyc2 genes from the four reference Chlorobia, and the 
cyc2 genes summarized from reference genomes by He and colleagues, 2017 (see reference in 
main text)

2. cyc2_Fig1_masked.faa -- the same as 'cyc2_Fig1_unmasked.faa', except run through 
Gblocks to mask (delete) ambiguous regions of the sequence alignment (see Methods). This 
was used to build the cyc2 phylogenetic tree (see Methods).

3. c5_family_Chlorobia_aligned.faa -- amino acid sequence alignment of the cytochrome c5 
family genes recovered from Chlorobia in this study, the cytochrome c5 family genes from 
the four reference Chlorobia.

4. cyc2_Fig3_other_hits_L227_L442.faa -- amino acid sequence alignment of the cyc2 genes 
recovered from non-Chlorobia genome bins from the metagenome datasets, as presented in 
Fig. 3B.
