## Supplementary File 7 for "Genomic potential for photoferrotrophy in a seasonally anoxic Boreal Shield lake": ABOUT_File_S7.txt

# ABOUT Supplementary File 7

This ZIP folder contains two files:

1. Ca_Chl_canadense_S13.3a_wet_mount_autofluorescence.png

Wet mount confocal laser scanning microscopy image of the "Ca. Chl. canadense" enrichment 
culture. The culture shown was inoculated into sulfide-containing Pfennig's medium using 
the same inoculum as for the photoferrotrophy test (Fig. 4) to serve as a positive 
control. Blue colour in the image represents autofluorescence at 600-800 nm wavelengths 
(presumably from bacteriochlorophyll c/d/e-containing cells of Chlorobia) when excited 
using a 488 nm laser, and grey colour represents transmitted light from the 488 laser. The 
scale bar represents 10 μm.

2. Ca_Chl_canadense_S11c_ASV_table.csv

Comma-separated table summarizing results from 16S rRNA gene amplicon sequencing (V4-V5 
region) of genomic DNA extracted from the "Ca. Chl. canadense" enrichment culture. The 
percent relative abundance of each amplicon sequencing variant (ASV) in the enrichment 
culture is shown, along with the taxonomy assigned to each ASV using the Silva database 
and the sequence of each ASV. Although this older subculture had evidently been overtaken 
by a heterotrophic contaminant (ASV1), the version of the culture used to test ferrous 
iron oxidation was dominated by phototrophic cells (see file #1 above).
